# Development of the entorhinal cortex occurs via parallel lamination during neurogenesis

**DOI:** 10.1101/2021.02.05.429755

**Authors:** Yong Liu, Tobias Bergmann, Yuki Mori, Juan Miguel Peralvo Vidal, Maria Pihl, Navneet A Vasistha, Preben Dybdahl Thomsen, Stefan E Seemann, Jan Gorodkin, Poul Hyttel, Konstantin Khodosevich, Menno P Witter, Vanessa Jane Hall

**Author notes:** Correspondence: Vanessa Hall.

## Abstract

The entorhinal cortex (EC) is the spatial processing center of the brain and structurally is an interface between the three layered paleocortex and six layered neocortex, known as the periarchicortex. Limited studies indicate peculiarities in the formation of the EC such as early emergence of cells in layers (L) II and late deposition of LIII, as well as divergence in the timing of maturation of cell types in the superficial layers. In this study, we examine developmental events in the entorhinal cortex using an understudied model in neuroanatomy and development, the pig and supplement the research with BrdU labeling in the developing mouse EC. We determine the pig serves as an excellent anatomical model for studying human neurogenesis, given its long gestational length, presence of a moderate sized outer subventricular zone and early cessation of neurogenesis during gestation. Immunohistochemistry identified prominent clusters of OLIG2+ oligoprogenitor-like cells in the superficial layers of the lateral EC (LEC) that are sparser in the medial EC (MEC). These are first detected in the subplate during the early second semester. MRI analyses reveal an acceleration of EC growth at the end of the second trimester. BrdU labeling of the developing MEC, shows the deeper layers form first and prior to the superficial layers, but the LV/VI emerges in parallel and the LII/III emerges later, but also in parallel. We coin this lamination pattern parallel lamination. The early-born Reln+ stellate cells in the superficial layers express the classic LV marker, Bcl11b (Ctip2) and arise from a common progenitor that forms the late deep layer LV neurons. In summary, we characterize the developing EC in a novel animal model and outline in detail the formation of the EC. We further provide insight into how the periarchicortex forms in the brain, which differs remarkably to the inside-out lamination of the neocortex.

## Introduction

The pig is an untraditional model in developmental neuroscience, but there are good reasons that warrant the use of the pig as a model, given its remarkable similarities to human neurodevelopment. Porcine fetal brain tissue can be easily obtained at specific time points of gestation when using controlled insemination procedures, which can be facilitated at pig production farms or in other controlled settings. Anatomically, the pig develops a gyrencephalic brain similar to humans, midway through gestation. Atlases (both annotated, unannotated and MRI) of the developing and adult pig brain are also readily available and accessible from open sources (Saikali, Meurice et al. 2010, Winter, Dorner et al. 2011, Conrad, Dilger et al. 2012, Conrad, Sutton et al. 2014). The pig brain increases significantly in size following birth, which is also reported in humans (Conrad, Dilger et al. 2012). Furthermore, there is limited proliferation of neurons in the postnatal large domestic pig brain, which is also observed in the postnatal human brain (Jelsing, Nielsen et al. 2006). Pig gestational development is relatively long, at 114 days, which is more comparable to human gestational length (294 days) than mouse gestational length (20 days). Given the reported divergence in cortex functions between rodents and humans (Farr, Kitas et al. 1988) and variations in the connectivity of the sensory cortical networks in the hippocampal region (Bergmann, Zur et al. 2016), it is necessary to consider other mammalian models which are more similar to human. To sum, the developing pig brain, albeit understudied in neurodevelopment, constitutes anatomically, a particularly useful model for modelling the developing human brain.

The structure of the entorhinal cortex (EC) in the fully formed brain presents an interesting interface between the 3 layered paleocortex and 6 layered neocortex, called the periarchicortex. The EC layer (L)IV in the EC is cell sparse and is better known as the *lamina dissecans*. (Insausti, Munoz-Lopez et al. 2017) The cell sparse LIV is a feature seen in all the allocortex subregions, including the paleocortex, archicortex and periallocortex (Stephan and Andy 1970, Stephan 1975, Stephan 1983). Whilst the paleocortex is comprised of 3 granular layers, the archicortex and the periallocortex (the latter is further subdivided into the periarchicortex and peripaleocortex) contains 4 granular layers (Stephan and Andy 1970, Stephan 1975, Stephan 1983). The EC has been subdivided into several areas but is now commonly divided into two, the lateral entorhinal cortex (LEC) and the medial entorhinal cortex (MEC) which deviates in both connectivity and function (Witter, Doan et al. 2017). The MEC is predominantly involved in spatial processing, whereas the LEC is predominantly involved in object recognition, odor discrimination and episodic memory (Staubli, Ivy et al. 1984, Schultz, Sommer et al. 2015, Vandrey, Garden et al. 2020). In the human, the EC is located in the anterior portion of the medial temporal lobe (Insausti, Munoz-Lopez et al. 2017), whereas, in the rat, it forms the ventro-caudal part of the cerebral hemisphere (Insausti, Herrero et al. 1997). Its location in annotated pig atlases indicates it lies in the piriform lobe, although the borders and boundaries have not been well annotated. The cell types within both the MEC and LEC are well characterized anatomically in other species and many cells have been profiled by electrophysiology (Canto and Witter 2012, Canto and Witter 2012). The principal neurons that reside in LII have received most attention. The LII comprises clustered pyramidal neurons expressing Calbindin (Calb) and scattered stellate cells in the MEC and fan cells in the LEC, both expressing Reelin (Reln) with subpopulations of pyramidal neurons, stellate cells, fan cells and multipolar neurons expressing both Calb and Reln (Witter, Doan et al. 2017). The stellate cells and pyramidal neurons in MEC LII contribute to the spatially modulated group of grid cells (Domnisoru, Kinkhabwala et al. 2013, Schmidt-Hieber and Hausser 2013, Rowland, Obenhaus et al. 2018), whereas the fan cells in the LEC are important for object-place-context associations (Vandrey, Garden et al. 2020).

The developing entorhinal cortex has only been studied to a limited extent. The medial EC is structurally organized already by embryonic day (E)18 in the rat (Bayer 1980, Bayer 1980, Bayer, Altman et al. 1993) and has formed, at least rostromedially, by gestational week (GW) 13-13.5 in the human (Kostovic, Petanjek et al. 1993). The maturation of the MEC has been studied during gestation and postnatal development via assessment in changes of molecular markers such as Calb, Doublecortin and Wolframin across the developing MEC (Ray and Brecht 2016, Donato, Jacobsen et al. 2017). The findings from these studies suggest the MEC circuitry matures along the dorsal to ventral axis, at least in the superficial layers. The stellate cells are one of the first cell types to be born and their maturation drives the maturation of the hippocampal and entorhinal circuitry ending in maturation of LII in the LEC (Ray and Brecht 2016, Donato, Jacobsen et al. 2017). Another study has characterized the expression of a few neuronal markers across the layers of the human EC at GW 20-26 (Zykin, Moiseenko et al. 2018), identifying Reln+ Cajal Retzius cells in LI but no Reln expression in the LII neurons. Differences were also observed in MAP2 expression. MAP2+ neurons could be clearly identified in only LIII of the LEC, but not in the MEC and LV of both the MEC and LEC. A more thorough investigation of neuronal markers is required during EC development as it is not clear, for example, if Reln is switched on in stellate cells at a later time point and what other markers can be used to delineate the different layers in the developing EC.

Evidence suggests that the EC may not form in the traditional inside-out lamination pattern as the rest of the neocortex. For example, an early study revealed through thymidine autoradiography in the developing rat brain, that LII formed earlier than LIII (Bayer, Altman et al. 1993). This has to some extent been confirmed by a more recent study by Donato et al., (Donato, Jacobsen et al. 2017) showing the birth of stellate cells early on during EC neurogenesis in mice. However, a detailed analysis of the lamination events during neurogenesis in the EC is lacking. Reln is thought to instigate a “detach and go” mechanism which instructs newborn neurons to detach from radial glia and signals them to translocate to the top of the cortical plate (Cooper 2008). Given Reln is found in both subsets of the LII and LIII neurons in the EC, it is unclear how this might affect cortical lamination of Reln-neurons in the same layers.

In this study, we sought to investigate the anatomical boundaries of the EC in the postnatal and developing EC in the pig to improve anatomical knowledge in a species that is understudied in neuroscience. We characterize the development of the EC using a large number of neural markers and investigate the pattern of cortical lamination in the EC, which forms part of the periallocortex, that differs in layer structure to the neocortex. Finally, we study the connectivity of the postnatal pig EC to elucidate its similarity to the more well studied rodents. We determine that the MEC develops by parallel lamination of the deep layers LV/VI followed by parallel lamination of the superficial layers LII/III.

## Materials and Methods

### Animal welfare and collection of brains

The experiments conform to the relevant regulatory standards for use of fetal material. Crossbred male and female pig fetuses at E26, E33, E39, E50, E60, E70, E80 and E100 of development were obtained from inseminated Danish Landrace/Yorkshire sows with Duroc boar semen from a local pig production farm with D0 counted as day of insemination. Deceased postnatal pigs were obtained at P75 as a gift from Per Torp Sangild at the University of Copenhagen. Adult brains were obtained from sows killed for another study using an overdose of sodium phenobarbital by a professional issued with a license from the Danish Animal Experiment Inspectorate.

### Brain Fixation and storage

For the earlier time points up until E60, the entire fetus was fixed and the dorsal skull was opened to improve permeation of the fixative. For the later time points, the brains were removed from the skull. Fixation was performed using 4 % paraformaldehyde (PFA; Millipore Sigma) in PBS (Thermo Fisher Scientific) from 24 hrs to up to 2 weeks, dependent on the size of the fetus/brain. Fetuses and brains were then stored long-term at 4°C in 0.002% Sodium Azide (VWR - Bie**&**Berntsen) in PBS.

### Paraffin embedding and sectioning

Prior to dehydration, the brains were dissected to approximately 10 mm thickness. The brains were dehydrated by immersion into a sequential series of ethanol, estisol and liquid paraffin using a tissue Processor (Thermo Fisher Scientific Citadel 2000). Following dehydration, the tissues were embedded in liquid paraffin followed by cooling down on a cold stage. Five-micrometer (μM) thick sections were cut on a microtome (Leica SM2000R) and mounted onto SuperFrost slides (Thermo Fisher Scientific). The sections were dried at room temperature (RT) overnight (ON) and stored long term at 4° C.

### Cresyl Violet Staining

Paraffin embedded sections were deparaffinized for 45 min at 60°C and 2x 10 min in Xylene. The sections were sequentially rehydrated for 2x 5 min in 99% EtOH, 3 min in 96% EtOH, 3 min in 70% EtOH, rinsed in tap water and stained for 12 min in 37°C Cresyl Violet (Millipore Sigma). The stained sections were rinsed in distilled water and staining outside of perikarya was differentiated for 10 min in 96% EtOH with 0.02% glacial acetic acid (Millipore Sigma). The sections were subsequently dehydrated 2x 5 min in 99% EtOH and 2x 5 min in Xylene prior to coverslip mounting using DPX (Sigma-Aldrich). All images were acquired on an Axio Scan.Z1 (Zeiss) using automated brightfield imaging.

### Tissue-Tek OCT embedding and Cryosectioning

Fixed fetuses/brains were dehydrated in 30% sucrose (Sigma-Aldrich)/1x PBS (Thermo Fisher Scientific) solution (0.22 μm filtered) at 4°C for 48 hours (h). The brains were cut into small pieces of approximately 4 mm thickness. The brain tissues were immersed into Tissue-Tek OCT (Sakura) and mounted within plastic molds (Simport) by snap-freezing in N-hexane ((VWR) solution immersed within liquid nitrogen. The frozen tissue was stored at −80°C until use. Brain sections were cut at 30 μm thickness using a cryostat (Leica CM 1950) and mounted onto SuperFrost Plus slides (Thermo Fisher Scientific) and stored at −20°C.

### Immunohistochemistry

Paraffin embedded brain sections were deparaffinized for 45 min at 60°C in the oven and 2x 10 min in Xylene (VWR). Subsequently, the sections were sequentially rehydrated for 2x 5 min in 99% EtOH, 3 min in 96% EtOH, 3 min in 70% EtOH, 3 min in tap water, and washed 2x for 5 min in PBS. The sections were subsequently permeabilized for 30 min in 0.1% Triton-X (Millipore Sigma) in PBS at RT and washed 2x 5min in PBS before antigen retrieval in boiling citrate [0.01M, pH6] (Sigma-Aldrich) for 3x 5 min. The sections were washed once for 5 min in PBS and Lab Vision™ MultiVision Polymer Detection System (Thermo Fisher Scientific) was used according to manufacturer’s protocol with the following specifications: Anti-parvalbumin (Millipore Sigma) primary antibody was diluted in 1:500 in PBS and incubated at RT for 30 min. The sections were visualized with LVRed and LVBlue.

### Immunofluorescence

Cryosectioned brains were air-dried for 1hr at RT, followed by rehydration in PBS (Thermo Fisher Scientific) for 10 min. After antigen retrieval the sections were permeabilized in 0.25% Triton X-100/PBS (Sigma-Aldrich) for 10 minutes followed by washing in PBS. The brain tissue was blocked in the blocking buffer (10% Normal donkey serum (Biowest) 2% BSA (Sigma-Aldrich)) at RT for 1h. Sections were incubated with the diluted primary antibody (see below) ON at 4°C followed by washing in PBS 3 times for 5 mins each. The sections were then incubated in diluted secondary antibody for 2h at RT and washed in PBS for 3 times for 5 mins each. The sections were counterstained with 10 μg/ml Hoechst 33258 (Sigma-Aldrich) for 10 min followed by washing in PBS for 5 mins twice. Secondary antibodies were conjugated with either Alexa fluorophores, 488, 594 or 647.

### Image processing and quantification

For acquisition of Cresyl Violet images, a bright-field microscope, Leica DMR with camera Leica DFC490 and Zeiss Axio scan.Z1 were used to capture the images. For acquisition of immunohistochemistry images, a confocal microscope Leica TCS SPE was used. Confocal images were acquired using Leica LAS X. Images were optimized for brightness and contrast using Fiji-ImageJ. Statistical analysis was conducted with commercially available software- Prism 7.0 (GraphPad Software).

### Structural MRI

A 9.4T BioSpec 94/30 USR spectrometer (Bruker BioSpin, Ettlingen, Germany) equipped with a 240 mT/m gradient system was used to acquire anatomical images of postmortem porcine brains (E60, E70, E80, E100, P75). Prior to imaging, the PFA fixed brain were suspended into plastic containers filled with a proton-free perfluorinated susceptibility-matching fluid (Solvay Galden® HT-230). We used a 35-mm (for P75) and a 23-mm (for E60, E70, E80, and E100) inner diameter transmit/receive volume coil (Bruker). The MR system was interfaced to a console running ParaVision software 6.0.1 (Bruker BioSpin). The parameters used in the brain scans were optimized for gray/white matter contrast: T2-weighted 2D rapid acquisition with relaxation enhancement (RARE) pulse sequence with TR/TE = 20000/40 ms (before birth), 12000/60 ms (after birth), Rare factor = 8 (before birth), 16 (after birth), in-plane resolution = 100 μm × 100 μm, slice thickness = 200 μm (before birth), 500 μm (after birth). 3D fast low angle shot (3D-FLASH) pulse sequence with TR/TE = 30/4.6 ms, Flip angle = 10, NA = 16, spatial resolution = 47 μm × 47 μm × 70 μm. The total imaging time was 2 hours.

### Diffusion MRI tractography

*Ex vivo* diffusion tensor imaging (DTI) of P75 brain (n = 3) was obtained using a Stejskal-Tanner sequence (TR/TE = 3500/17.5ms, NA = 2, spatial resolution = 390 μm × 390 μm × 390 μm) on a 9.4T spectrometer equipped with a 1500 mT/m gradient system and a 40-mm inner diameter transmit/receive volume coil. We applied the motion probing gradients (MPGs) in 60 noncollinear directions with b = 2500 s/mm^2^ in addition to 4 b0. Diffusion Toolkit (version 0.6.4.1) and TrackVis (version 0.6.1) were used for 3D reconstruction of white matter tracts. We used a DWI mask threshold and an angular threshold of 45 degrees. Two regions of interest (ROIs) were selected a priori for the DTI analysis: including the LEC and MEC. ROIs were created by manually segmentation with ITK-snap (version 3.6.0, (Yushkevich, Piven et al. 2006)). The fiber tracks obtained from the ROIs were qualitatively analyzed for the direction of tracks, and differences in FA values within the brain parenchyma. The assessment was done using tractography maps: (1) a standard color-coded reconstruction to visualize the location and orientation of white matter pathways (blue tracks represented the axonal orientations in the craniocaudal, green in anterior-posterior, and red in left-right direction); (2) a scalar FA map with minimum and maximum FA thresholds of 0.1 and 0.6, respectively; and (3) total track length information for each ROI.

### BrdU labeling

Mouse experiments were carried out in accordance with the guidelines of the National Animal Ethics Committee of Denmark. In the present study, C57BL/6 wild-type mice (Janvier Labs) were used. The mice were housed in individually ventilated cages in a reversed light cycle and with free access to food and water. Males and females were housed together until a vaginal plug (VP) was detected, after which the males were separated. Detection of the VP, was checked each morning, and if observed was then defined to be E0.5. Neuronal progenitors were labelled with bromodeoxyuridine (BrdU, Sigma-Aldrich B5002) by intraperitoneal injection (i.p) of 50 mg/kg BrdU into dams at E8.5 to E16, with half-day intervals (with the exception of E14.5 and E15.5). Note, each mouse received only one injection. The concentration was previously validated to label progenitor proliferation in the embryonic brain without inducing obvious toxic effects (Vasistha, Johnstone et al. 2019). Pups were anaesthetized by i.p. injection with a combination of xylazine and ketamine on postnatal day (P)7 and perfused transcardially, first with 10 mL cold PBS and subsequently with 5 mL cold 4% PFA. The brains were removed and fixed overnight (O/N) in 4% PFA at 4 °C and stored in PBS with 0.02% sodium azide.

### Cryo-embedding and sectioning of mouse brains

Following PFA fixation, mice brains were dehydrated in 30% sucrose (Sigma-Aldrich) in PBS O/N at 4 °C. One hemisphere was embedded by immersion in O.C.T. Compound (Tissue-Tek, Sakura) in a cryomold, snap-frozen in N-hexane (VWR) submerged in liquid nitrogen, and stored at −20 °C O/N until used for cryosectioning. The brains were sectioned on a cryostat (Leica CM 1950) at 30 μm thickness and mounted onto SuperFrost Plus slides (Thermo Fisher Scientific) and stored at−20 °C.

### Statistics

For immunohistochemistry and assessment of neuroconnectivity from DTI white matter tracts, all samples were performed in biological triplicates. Statistical analysis was conducted using commercially available software- Prism 7.0 (GraphPad Software). Dependent on the experimental design, an ordinary one-way ANOVA or an unpaired two-tailed t test was performed to statistically assess differences. All error bars denote standard deviation (SD). Non-significant p-value (n.s.p) >0.05. Significant differences are marked in figures using asterixes which represent the following: * P≤0.05, ** P≤0.01, *** P≤0.001.

The counting of NeuN and BrdU immunolabelled cells in the mouse brain sections was performed manually using the Cell Counter tool in Fiji. The cell counting was performed on individual z-plane images that were later merged, to ensure that overlapping cells were counted only once. Anti-BrdU labeled cells were only counted as positive if a strong labelling was observed (at least 50% staining). The quantity of NeuN and BrdU positive cells were counted in the layer (L)II, LIII and LV / VI of the MEC in sections from three separate brains with BrdU injection at each time point. For the remaining stages of BrdU injection only two separate brains were quantified. The fraction of neurons with BrdU incorporation was calculated as the fraction of BrdU positive/NeuN positive cells. The mean and SD of the fraction of BrdU positive neurons was calculated in R for the different layers at each individual stage of BrdU injection. The SD was calculated in R, using the sd() function.

## Results

### Subheading 1. The MEC and LEC boundaries can be delineated in the prenatal porcine brain

The pig is not traditionally used as a model to study the EC or brain development, hence, it was important to validate that the pig EC was highly similar to other species. Gestational age differences and trimester lengths were first compared between mouse, man and pig to determine when corticogenesis occurs in the pig. The transition from trimester 1 to 2 is an important early time point, since this is when neural tube formation and gastrulation is completed in mouse and man (Patten, Fontaine et al. 2014). Other key developmental features, such as fusion of the palate also mark this transition, as well as gonadal differentiation. Gonadal differentiation has been studied in the pig and occurs at E (embryonic day) 50 in the pig (Pontelo, Miranda et al. 2018). We therefore deduced that the end of the first trimester occurs at E50. The transition from trimester 2 to 3 marks a significant and continual increase in human fetal weight upon completion of organogenesis. A similar and marked increase in fetal weight occurs at E70 in the pig (Kim 2010). We believe, therefore, that E70 marks the transition from trimester 2 to 3. We highlight this comparative overview of trimester lengths between species in Figure 1A.

**Figure 1.**
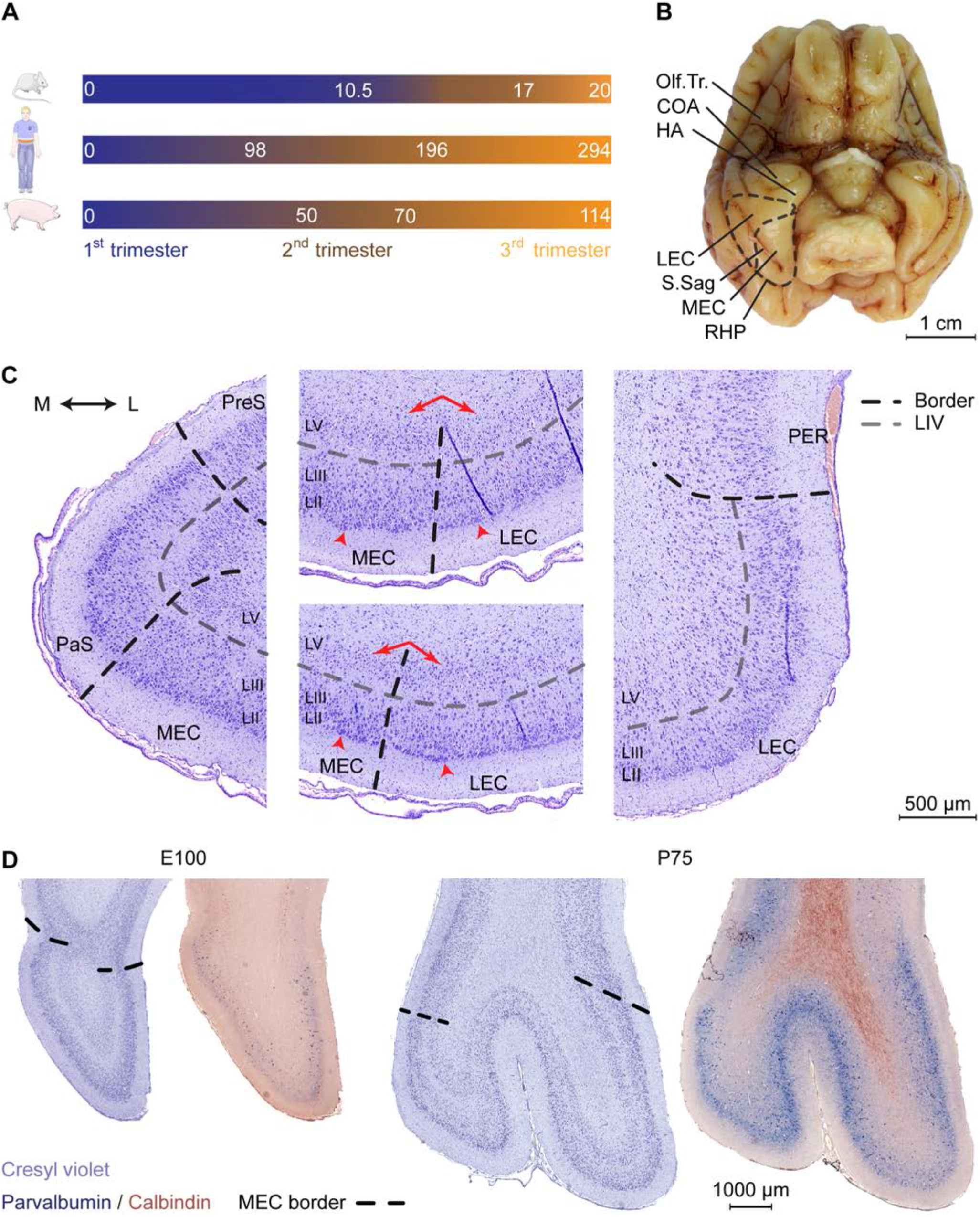
Cytoarchitectonic features of the developing porcine entorhinal cortex. **(A)** Length and trimester divisions of the gestation in mouse, human, and pig. **(B)** Macroscopic view of the ventral surface of the porcine brain at Embryonic day (E)100 with annotation of the piriform lobe including the entorhinal area. Dashed black line indicates the outline of EC and the border between lateral and medial entorhinal cortex. Olfactory tract (Olf.Tr.); caudate nucleus of amygdala (COA); hippocampal area (HA); lateral entorhinal cortex (LEC); medial entorhinal cortex (MEC); sagittal sulcus (S.Sag); posterior rhinal sulcus (RHP; note that the sulcus coincides with the visible lateral border of the EC). **(C)** Delineations and cytoarchitectonic features of cresyl violet stained coronal sections of the EC (shown at E100 as representative). Light grey punctuated line marks the acellular layer (L)IV l*amina dissecans*, black punctuated line borders between MEC and LEC or EC and adjacent areas. Red arrows (middle panels) shows the difference in the cellular organization in deep layers, arrowheads highlight large superficial cells of the EC. Medial (M); lateral (L); pre-subiculum (PreS); para-subiculum (PaS), perirhinal cortex (PER). **(D)** Cresyl violet staining and parvalbumin (PVALB) expression highlights the caudal boundaries of the EC at E100 and P75 (coronal plane). PARV expression coincides with defined cytoarchitectonic borders of the EC. Scale bar 1000 μm.

We then performed a detailed characterization of the location of the EC in a late gestational pig brain at E100 (2 weeks prior to birth). In the pig, the EC is located in the piriform lobe, caudal to the olfactory (piriform) cortex and the cortical nucleus of amygdala (COA) and hippocampal transition area (HA) (Figure 1B). Cresyl violet staining revealed a cytoarchitecturally mature EC, with similar morphology as the postnatal EC (Figure 1C and Figure S1). Layer IV was distinctively acellular, which is a typical feature in the adult EC known as the *lamina dissecans*. We identified the parasubiculum (PaS) medial to the EC. The PaS could be identified at E100 from its similar cytoarchitecture as the EC. One exception was that the soma sizes of the PaS LII and LIII cells were more equal, compared to the EC. In the EC, the LII cells have somas approximately twice as large as in the LIII (Witter, Doan et al. 2017). We could identify the perirhinal cortex (PER) located directly adjacent and lateral to the EC through the disappearance of large superficial cells of the LII typical of the EC, together with the disappearance of the *lamina dissecans* (Figure 1C). Already at E100, the EC can be easily divided into a lateral (LEC) and medial (MEC) subdivision. This is based on cytoarchitecture differences described in other species such as rodents, dogs, cats and humans (Wyss, Sripanidkulchai et al. 1983, Insausti, Tunon et al. 1995, Woznicka, Malinowska et al. 2006, Gatome, Slomianka et al. 2010). The pig MEC presented deeper layers structured with a narrow homogenous and densely packed cellular layer organized with a columnar appearance. In the transition into the LEC, the deep layers widened with a lower cell density and a disordered cell deposition. The LIV *lamina dissecans* in the MEC was distinct and acellular, while in the LEC, the *lamina dissecans* was more diffuse, with more cellular infiltration (Figure 1C). The MEC gradually occupied the entire mediolateral entity along the rostrocaudal axis of the piriform lobe. The MEC caudally borders to the neocortex as the piriform lobe disappears. This boundary was traced by immunolabeling for parvalbumin (PVALB) positive interneurons (Figure 1D), as seen in the rat (Wouterlood, Hartig et al. 1995). We observed a sulcus forming at the mid-rostrocaudal extent of the EC, persisting to the caudal end of the piriform lobe which was also visible macroscopically (Figure 1B). This sulcus, known as the sagittal sulcus (S.Sag) has been previously reported in the domestic (Holm and Geneser 1989) and Göttingen minipig (Bjarkam, Glud et al. 2017), but not in other gyrencephalic species, such as dogs, cats, and humans (Wyss, Sripanidkulchai et al. 1983, Insausti, Tunon et al. 1995, Woznicka, Malinowska et al. 2006). We observed the S.Sag to coincide with the appearance with the MEC on the rostrocaudal axis (Figure S1), which provided a unique surface landmark for microscopically and macroscopically assessing the MEC and LEC borders. This anatomical study was important for assessing the borders of the MEC for further detailed investigation.

### Subheading 2. Prominent oligodendrocyte precursor cells are a major anatomical feature in the superficial layers of the developing porcine LEC and perirhinal cortex

Interestingly, we observed a prominent number of oligodendroglia-like cells present in the superficial layers of the LEC and in the adjacent perirhinal cortex (PER), which has not been previously highlighted in other anatomical descriptions of the EC. These cells were proximal and clustered around large entorhinal cells (Figure 2A). Morphologically, these appeared to be pan-neuronal oligodendrocyte precursor cells (OPCs) with clear Nissl free cytoplasm and a large round Nissl stained nucleus (Garcia-Cabezas, John et al. 2016) (Figure 2B). We observed an enrichment of these oligodendroglia-like cells in the superficial layers of the LEC and PER from E80 of development and fewer cells were observed in the MEC and cingulate gyrus (dorsal neocortex) (Figure 2B). In the most lateral part of the LEC, these OPC-like cells were evenly distributed within the superficial layer but converged into distinct islets towards the MEC. The islets disappeared at the border of the LEC-MEC and the OPC-like cells were more evenly distributed within the MEC. We were not able to distinguish these cells using cresyl violet staining prior to E80 (Figure S2), therefore we immunolabeled the MEC from E50 onwards with the OPC marker, OLIG2 in order to determine if they may be present earlier. We detected OPC+ cells from E50 onwards (Figure 2C). At E50, the OLIG2+ cells were detected predominantly in the subplate and from E60 onwards were distributed more evenly across the developing cortical layers (Figure 2D).

**Figure 2.**
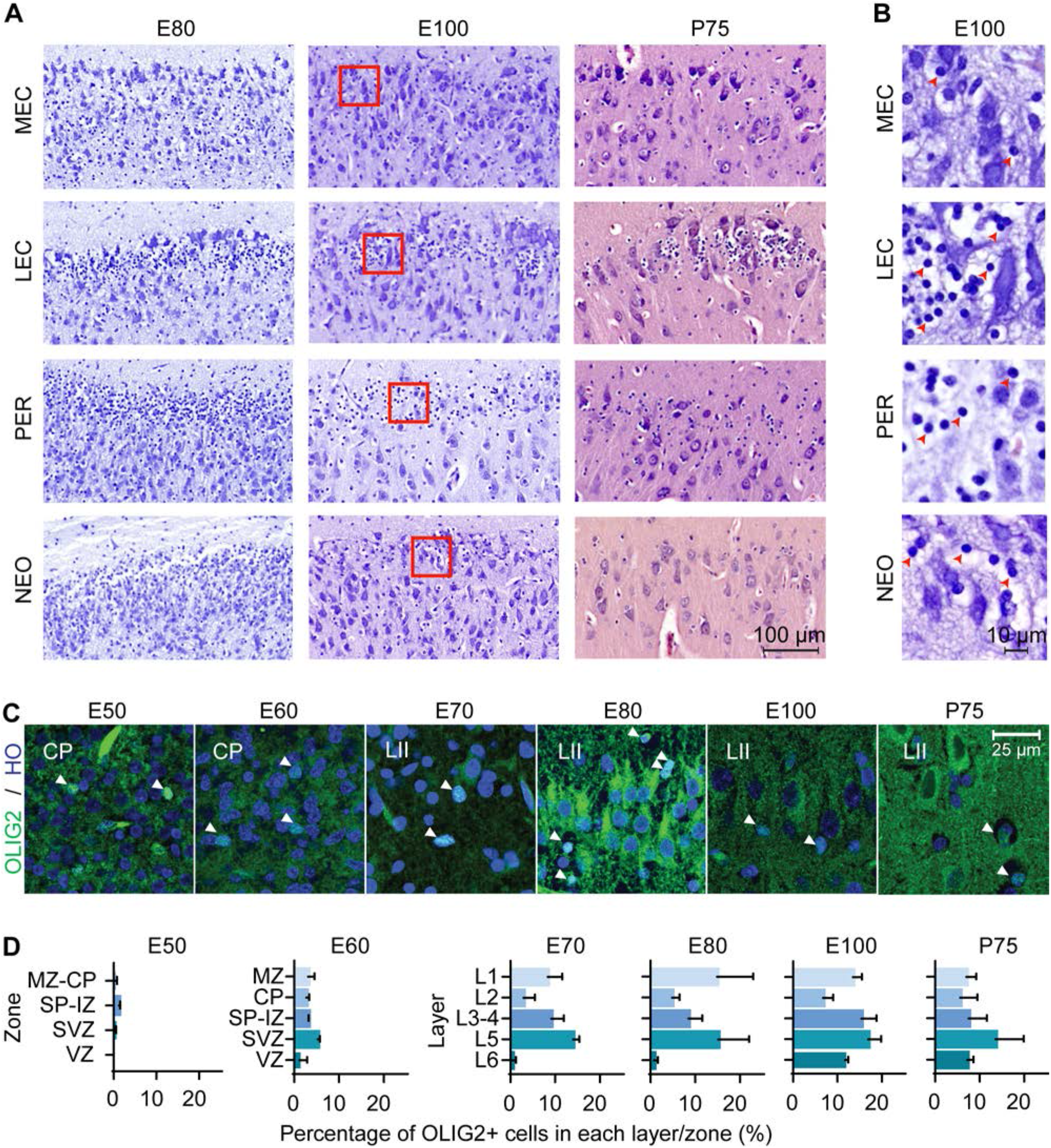
Oligodendrocyte precursor cells (OPCs) emerge in the entorhinal cortex (EC) early in the second trimester. **(A)** A high number of OPCs are present in the superficial layers of the lateral entorhinal cortex (LEC) and perhirhinal cortex (PER) and less OPCs observed in the medial entorhinal cortex (MEC) and neocortex (NEO). Scale bar 100 μm. **(B)** High magnification of red squares in (A) depicts the morphology of the OPCs at Embryonic day (E)100 which shows condensed nuclei and nissl free cytoplasm (red arrowheads). Scale bar 10 μm. **(C)** OLIG2 expression and Hoescht (HO) staining in the developing EC. Images taken within the developing cortical plate (CP) (E50-E60) and later (E70-P75), in Layer II. Scale bar 25 μm. **(D)** Quantification of OLIG2 positive cells in the EC by zones/cortical layers from E50 until postnatal day (P)75. Cortical plate, CP; intermediate zone, IZ; marginal zone, MZ; subplate, SP; subventricular zone, SVZ ventricular zone, VZ. All error bars represent SD.

### Subheading 3: The developing EC accelerates in growth at the end of the second trimester

We subsequently analyzed the developmental timing of the EC using cresyl violet staining (E50-P75) and structural postmortem MRI (E60-P75). No features of an EC could be determined prior to E50. At E50, we identified prominent and large soma of entorhinal-like cells in the superficial layer of the ventral telencephalon before any distinct layer was formed (Figure 3A). The three-layered cortex became six-layered by E60 and the *lamina dissecans* was now visible (Figure 3B). At E60, we were able to delineate the MEC from the LEC based on the deposition of the deep layer cells and the higher cellularity of the *lamina dissecans* in the LEC. Together, this indicated that the EC cortical layers form during the second trimester between E50 and E60 in the pig. At E70 (late in the second trimester), we observed the S.Sag formation microscopically and cells in the superficial LII were prominent, large and darkly stained nuclei with Nissl (Figure 3C). From E80, the large cells in LII appeared more prominent (Figure 3D) and the S.Sag was macroscopically visible (Figure 3E).

**Figure 3.**
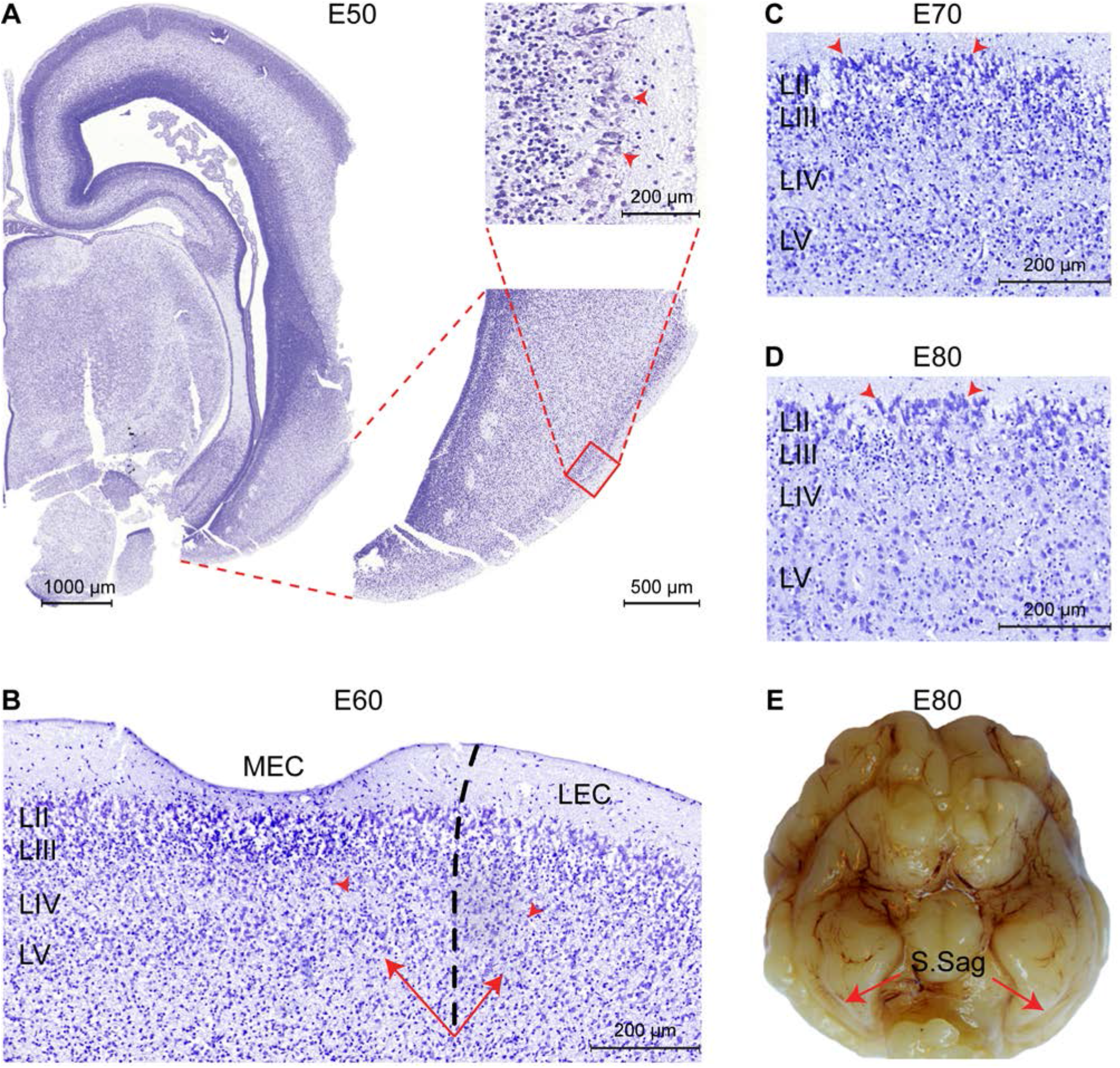
The entorhinal cortex (EC) forms between Embryonic day (E)50 and E60 in the pig. **(A)** Entorhinal-like cells with large somas (red arrowheads) are identifiable at embryonic day (E)50 in the superficial layer of the ventral telencephalon. **(B)** At E60 the EC becomes six-layered, the *lamina dissecans* becomes apparent as layer (L)VI, and the medial EC (MEC) and lateral EC (LEC) subfields of the EC become distinguishable, based on the difference in cellularity of the lamina dissecans (red arrowheads) and the deep layer (LV-LVI) organization (arrows). Scale bar 500 μm. Section of the MEC at E70. Prominent, large and darkly stained nuclei in the superficial LII (arrowheads) are apparent. Scale bar 200 μm. **(D)** Section of the MEC at E80. Superficial LII cells become more prominent. Scale bar 200 μm. **(E)** The sagittal sulcus (S.Sag) becomes macroscopically visible in the piriform lobe at E80 (red arrows).

The positional anatomy and rate of EC growth was evaluated using postmortem structural-MRI. The EC was annotated from E60 to P75 based on our histological descriptions and macroscopic features (Figure 4A). We calculated the volume of the EC and the remaining cortex based on the MRI annotations and found the EC growth was linear from E60 to P75 (Figure 4B). When we compared the growth rate of the EC to that of the whole cortex of the brain, we found that the EC had a significantly higher growth rate at E70 during the late second trimester (Figure 4B), suggesting a local and specific developmental growth period at this time point. We were unable to detect whether the growth was due to expansion of the white or gray matter, due to the lack of MRI resolution. In summary, we identified a significant growth spurt during the late second trimester attributable to either grey and/or white matter expansion.

**Figure 4.**
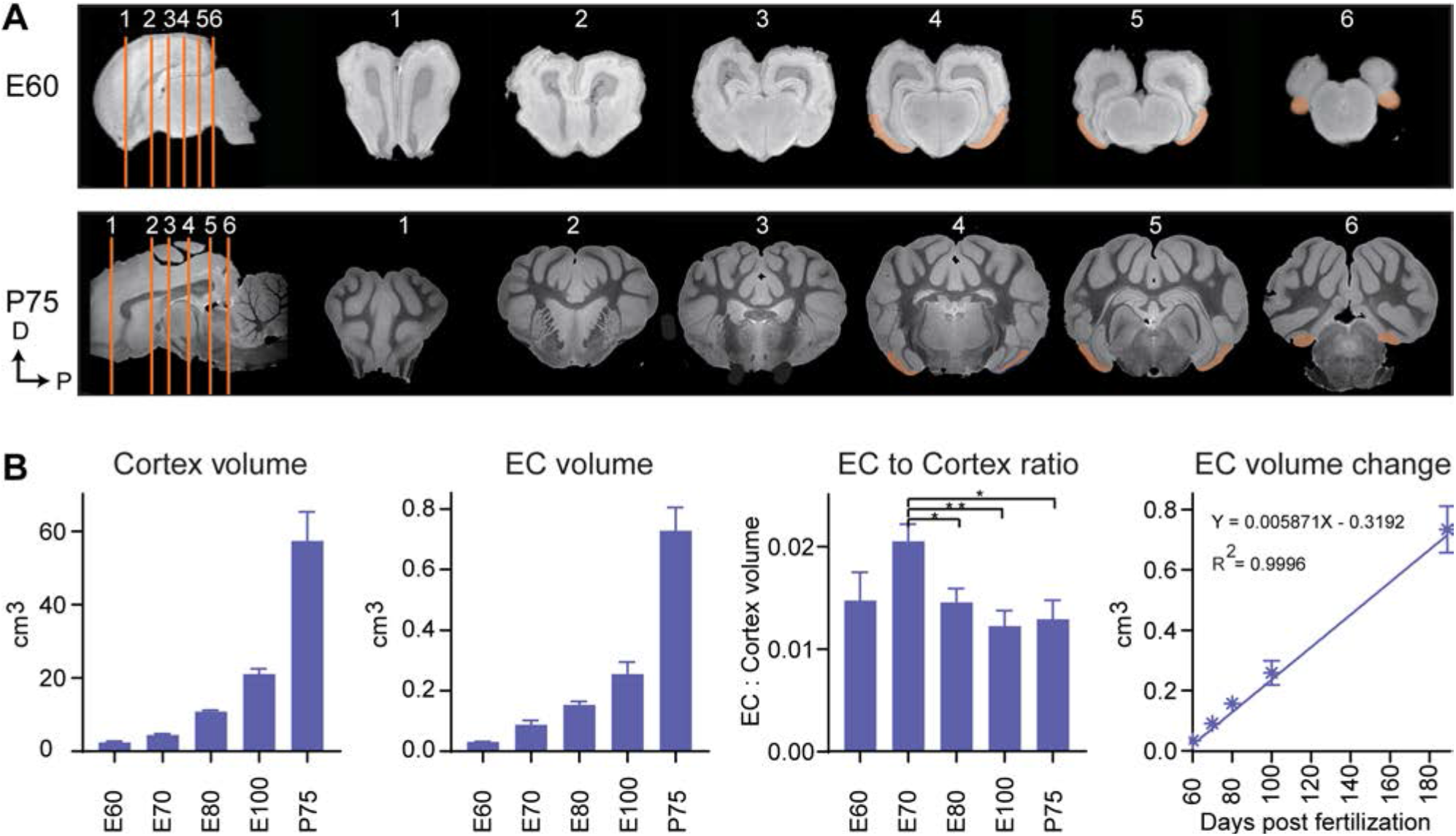
Magnetic resonance imaging (MRI) of the developing entorhinal cortex (EC) shows extensive growth at the end of the 2^nd^ trimester. **(A)** MRI shows the rostral to caudal position of the early developing EC (embryonic day (E)60) and postnatal EC (postnatal day (P)75) in the coronal plane (sections depicted by orange lines). **(B)** The total cortex volume, the EC volume, the EC:Cortex ratio and the change in EC volume over time shows an overall linear growth pattern during development, but an exceptional growth spurt of the EC compared to the rest of the neocortex at E70 during development.

### Subheading 4: Neuronal projections from the postnatal EC reach several brain regions

We then analyzed the EC and its connectivity to other brain regions using DTI tractography in a later and more developed EC, in postnatal (P) brains 75 days after birth (P75). A high number of white matter tracts emerged from both the MEC (median 6031±1319) and the LEC (median 3329±4099) (Figure 5A, B, Supplementary Table 1). Projections extended to the peri- and post-rhinal cortex, the hippocampus, hippocampal commissure, olfactory bulb, amygdala, nucleus of the lateral olfactory tract, putamen and nucleus accumbens (Figure 5C-H, Supp video 1, Supp Table 1). Further connections between the MEC and LEC could also be observed (Supp Table 1, Supp video 1). The MEC connected to more regions than the LEC (Figure 5H) and most connections were observed to the amygdala and subiculum (Figure 5H, Supp Table 1). Most of the LEC connections traced also to the amygdala (Figure 5H, Supp Table 1). Regarding connectivity to the hippocampus, both the LEC and MEC connected to the DG, CA1 and CA3 regions, although more connections were captured from the MEC (Figure 5H, Supp Table 1). The observed connectivity to the peri- and post-rhinal cortex concurs with previous studies reported in the rat (Insausti, Herrero et al. 1997, Burwell and Amaral 1998). A small number of fibers that emerged from the MEC projected to the dorsal hippocampal commissure (Figure 5G, Supp table 1). Strikingly, most of the tracts traced to other regions with a large number extending to the rostroventral brain, presumed to be the orbitofrontal cortex or potentially the anterior visual cortex (which remains unannotated in the pig). Many LEC fibers ending at the rostral end of the brain, ended in a slightly caudal (unannotated) location to the MEC fibers (Supp Video 1). To sum, the connectivity between the EC and septohippocampal region in the postnatal pig brain was remarkably similar to that which has previously been reported in other species, including humans (Wilhite, Teyler et al. 1986, Witter, Room et al. 1986, Mufson, Brady et al. 1990, Totterdell and Meredith 1997, Kolenkiewicz, Robak et al. 2009, Rowland, Weible et al. 2013, Sun, Nguyen et al. 2014, Witter, Doan et al. 2017) which helps to consolidate it as an excellent model for human EC connectivity.

**Figure 5.**
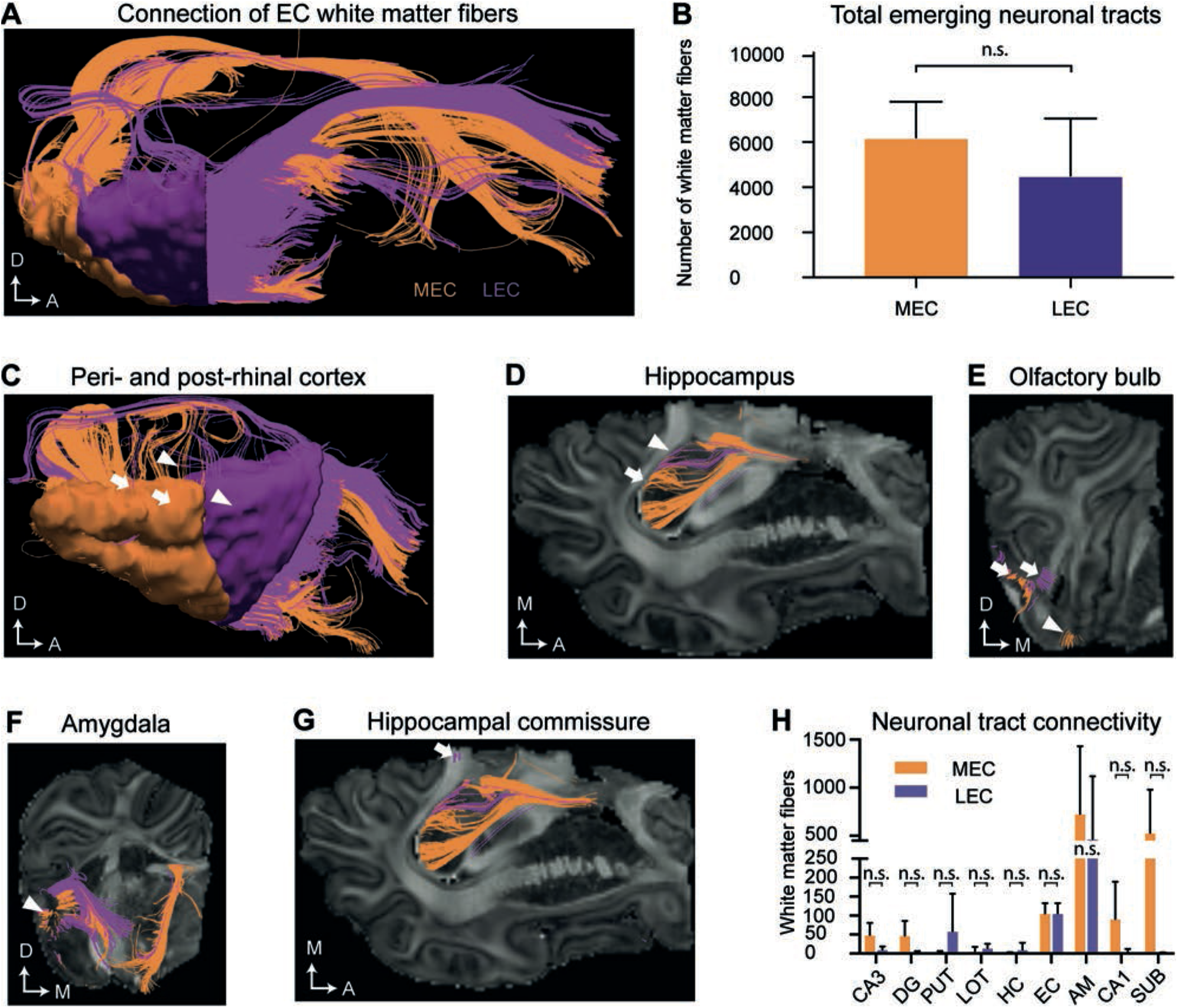
Diffusion tensor imaging (DTI) of the white matter projections identify white matter fiber projections to varying brain regions at P75. **(A)** An overview of white matter fibers connecting to/from the medial entorhinal cortex MEC (orange) and laternal entorhinal cortex LEC (purple). **(B)** The average number of tracts connecting to/from the MEC and LEC do not differ significantly. **(C)** White matter fibers connecting to/from the perirhinal cortex (white triangles) and postrhinal cortex (white arrows). **(D)** White matter fibers connecting to the hippocampus (white arrows). **(E)** White matter fibers connecting to the lateral (white arrows) and medial olfactory bulb (white triangle). **(F)** White matter fibers connecting to the amygdala (white triangle). **(G)** White matter fibers extend from the EC to the hippocampal commissure (white arrow). Axes: A, Anterior; D, Dorsal, M, Medial. **(H)** The average number of tracts connecting to different anatomical regions emerging from the EC do not significantly differ between the MEC and LEC. All error bars denote SD. n.s, not significant. Significance threshold, p ≤ 0.05. Abbreviations, AM: amygdala; DG: dentate gyrus; EC: entorhinal cortex; HC, Hippocampal commissure; LOT, lateral olfactory tract; PUT, putamen; SUB: subiculum.

### Subheading 5. The developing EC is marked by parallel lamination

Given evidence suggests the layers of the EC may not form or mature in the classic inside-out lamination pattern, we decided to trace the cortical lamination patterning and focused on the MEC where the stellate cells reside and are known to drive maturation events (Donato, Jacobsen et al. 2017). The timing of neurogenesis was investigated by performing immunohistochemistry at 8 time points from E23 to P75 of development. Using antibodies directed against GFAP, FABP7 (BLBP), SOX2 and PAX6, we could detect the neural epithelium (NE) (PAX6+/FABP7-) at E23 and radial glia (GFAP+/FABP7+) at E26 (Figure S3A-C). We measured the size of the ventricular zone (VZ) (PAX6+) and identified that it was present at E23 and peaked in size at E50 (Figure S3D). After this time point, the VZ declined dramatically in size at E60 (Figure S3D). We then investigated the second germinal zone by performing immunohistochemistry using antibodies directed against EOMES and PAX6. Our results showed the presence of a subventricular zone (SVZ) containing different populations of EOMES/PAX6 progenitors (Figure S3E-F). Later in neurogenesis the SVZ splits into an inner- and outer-subventricular zone (ISVZ and OSVZ), with the OSVZ being much larger in size in humans and non-human primates, compared to ferrets and rats (Smart, Dehay et al. 2002, Fietz, Kelava et al. 2010, Martinez-Cerdeno, Cunningham et al. 2012). Assessment and quantification of the porcine SVZ at E26 showed that EOMES+/PAX6+ cells and EOMES+/PAX6-cells were present at an equal ratio (Figure S3F). At E33, two distinct layers could be observed above the VZ, a dense inner subventricular zone (ISVZ) containing equal proportions of EOMES+/PAX6+ and EOMES+/PAX6-cells and a diffuse overlying OSVZ containing EOMES+/PAX6+ and EOMES+/PAX6-cells (Figure S3E-F). We observed that the ratio between EOMES+/PAX6+ and EOMES+/PAX6-cells changed over time, with EOMES+/PAX6+ cells becoming more abundant in both the ISVZ and OSVZ (Figure S3F). These zones had declined by E70. Considering PAX6 is an important transcriptional regulator of neurogenesis (Sansom, Griffiths et al. 2009), our findings imply that neurogenesis within the porcine EC occurs from E33 until shortly after E60.

We then analyzed the events of cortical lamination in the EC by using immunohistochemical analyses of canonical markers for deep (BCL11B) and superficial layers (SATB2), as well as EOMES, RELN and TBR1 across varying times of gestation in the pig. During early cortical formation (E21), we detected the presence of EOMES+ and RELN+ cells towards the pial surface (Figure S3G). Co-expression of EOMES+/RELN+ cells were detected 2 days later at E23 and expression of TBR1 emerged in some of these cells at E23 with more RELN+/TBR1+ cells detectable by E33 (Figure S3G-H) which we deduced to be migrating and maturing Cajal–Retzius cells (CR cells) that are important for regulating cortical lamination (Hevner, Neogi et al. 2003). We then traced the emergence of local neurons using BCL11B and SATB2. As expected, BCL11B was the first of the two markers to be expressed at E33 and was located in the cortical plate (Figure S4). Interestingly, a large proportion of the BCL11B+ cells expressing SATB2 in the cytoplasm could be detected in the main cortical layer at E39 (Figure S4). By E50 a distinct new population of BCL11B cells had emerged as a superficial layer of the EC which could not be observed in the dorsal telencephalon (Figure 6A-B). These large prominent nuclei resembled the prominent large entorhinal cells that were observed histologically (Figure 1C). In the dorsal telencephalon, a SATB2+ population emerged in the superficial layer at E50 which remained present until gestation (Figure 6A-B) which was not detected in the EC. In contrast, a new population of BCL11B+ cells emerged at E60 underneath the superficial layer of BCL11B+ cells. This indicated that the porcine EC was forming in a non-conventional pattern. This cell layer positioning was retained even after birth (Figure 6C) and was not observed in the dorsal telencephalon (Figure 6B). A small proportion of the LII BCL11B+ cells began to express RELN at E60 (10 days after deposition) with expression becoming more abundant in the superficial cells at E70 (Figure 6B, C). The RELN+ neurons residing in both the MEC and LEC continued to express BCL11B and SATB2 after birth (Figure 6D, E).

**Figure 6.**
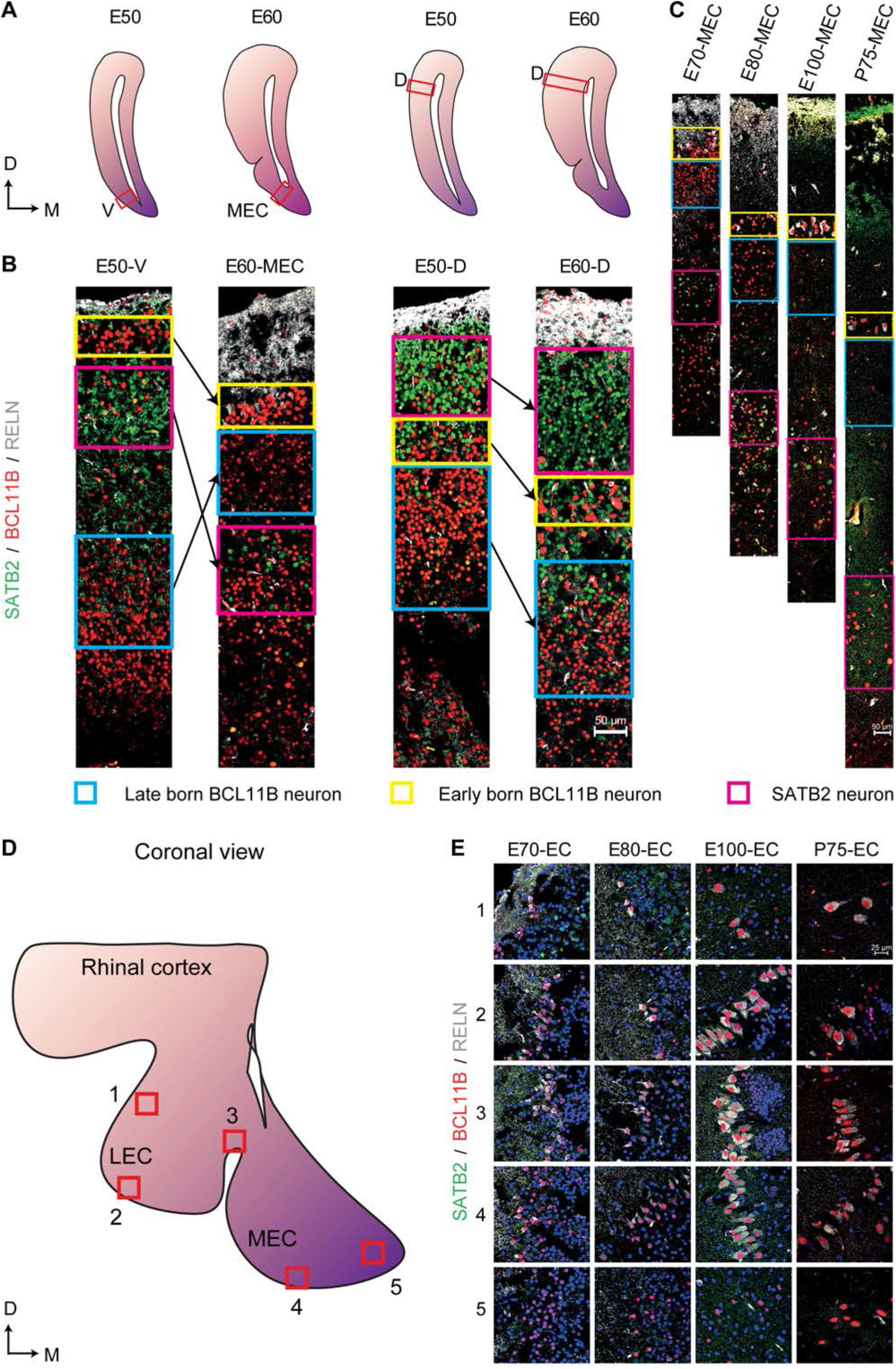
Altered expression of superficial and deep layer markers in the entorhinal cortex (EC) persist even after birth. **(A)** Locations of the analyzed ventral telencephalon (V), the MEC, and the dorsal telencephalon (D) at Embryonic day (50) and E60 of development (red boxes). **(B)** Expression of SATB2, BCL11B and RELN in the ventral telencephalon, within the MEC and the dorsal telencephalon at E50 and E60 shows a prominent expression of BCL11B in the superficial layer of the MEC at E50 and the emergence of a new BCL11B population beneath the BCL11B+ LII in the MEC at E60. **(C)** Expression of SATB2, BCL11B, and RELN in the MEC from E70 until Postnatal day (P)75 of development shows persistent expression of BCL11B+ cells in the superficial layers. Locations of analyzed superficial layers across the MEC and LEC from E70 to P75. **(E)** RELN+ neurons co-express BCL11B across the EC during gestation and also after birth. Scale bar 50 μm **(B, C)** and 25 μm **(E)**.

An independent approach that would support our observation of non-conventional pattern of MEC formation would be to study the timing of cortical layer deposition using bromodeoxyuridine (BrdU) labeling. Since the quantities of BrdU required for administration of >200 kg pregnant sows were not feasible, we selected the mouse as a translational model. Injections were performed at half day intervals from E8.5 to E14 and one day intervals from E14 to E16 to capture the entire span of neurogenesis. Brains were collected from P7 mice and BrdU labeling was only measured in strong labelled cells. We observed the birth of NeuN+ neurons simultaneously in the deeper layers, LV/LVI as early as E10.5 with a peak occurring at E12.5 (Figure 7A, Figure S5A-B). The birth of both LII and LIII NeuN+ neurons was observed later at E12.5, but interestingly, both layers were born in parallel and both peaking at E14 (Fig 7A, Figure S5A-B). The birth of LV/VI neurons was mostly complete by E15 and the birth of LII and LIII neurons was complete by E16 (Fig 7A, Figure S5A-B). Interestingly, at E12.5, the BrDU labeling was weaker in the superficial neurons suggesting that they formed after the birth of the deep layer neurons and likely arise from a common progenitor (Figure S5). Collectively, our findings show that LII and LIII neurons are born after the deeper layer neurons, however, the deeper layers LV/LVI arise in parallel and the superficial layers LII and LIII also arise in parallel as opposed to an inside-out deposition of layers. We therefore name this lamination process parallel lamination.

**Figure 7.**
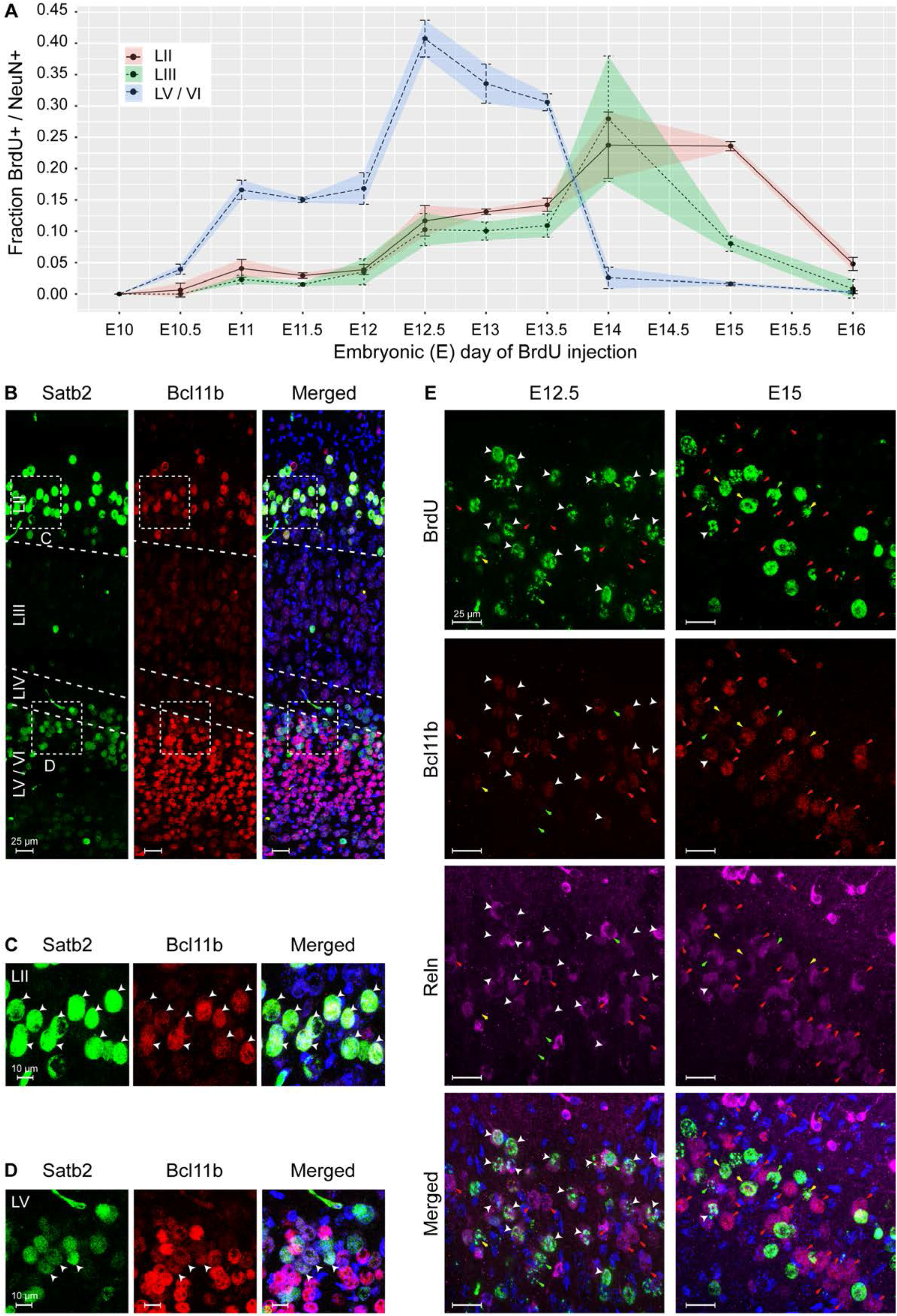
The medial entorhinal cortex (MEC) emerges via parallel lamination events. Birth timing of layers in the MEC was established using Bromodeoxyuridine (BrdU) labelling of NeuN-positive (+) neurons born at embryonic day (E)10 to E16 and identification of early-born neurons by expression of Satb2, Bcl11b, and Reln. **(A)** Quantification of NeuN+ neurons labelled with BrdU in LII, LIII and LVa/b/VI, respectively, at the time of BrdU injection, shown as correlation of BrdU+/NeuN+ fraction and injection time in gestation as embryonic day (E). Error bars denote the standard deviation. **(B)** Overview of Satb2 and Bcll1b expression in the medial entorhinal cortex (P7). The upper-layer (UL) marker, Satb2 was expressed in the LII and LVa. The deep-layer (DL) marker Bcl11b, was expressed both in the DL and co-expressed with Satb2 in neurons in LII. Merged image additionally shows Hoechst (HO) staining. Scale bar 25 μm. High magnification of UL **(C)** and DL **(D)** inserts from A, show Satb2+ and Bcl11 expression in LII cells. White arrow heads point to cells with co-expression. Merged image additionally shows HO DNA labeling. Scale bar 10 μm. **(E)** Co-expression of Bcl11b and Reln in cells in the LII newborn neurons at E12.5 and E15 (assessed by BrdU labeling). White arrow heads show cells with immunoreactivity for BrdU, Bcl11b and Reln. Red arrows point to cells co-expressing Bcl11b and Reln. Yellow arrows depict expression of Bcl11b in BrdU labelled cells. Green arrows depict expression of Reln in BrdU labelled cells. Merged image shows immunoreactivity of BrdU, Bcl11b, Reln, and HO. Scale bar 25 μm.

Similar to the pig, we found that postnatally in mice (P7), cells in both LII/LIII and LV co-expressed Bcl11b and Satb2 (Figure 7B-C). Interestingly, a mixture of populations was present in the LV, including Bcl11b+/Satb2-, Satb2+/Bcl11b- and Bcl11b+/Satb2+ cells (Figure 7B-C). LVI cells were Bcl11b+/Satb2-. We only observed a single population of Satb2+/Bcl11b+ cells in LII/LIII. To investigate the emergence of the Reln+ neurons in LII in more detail, we investigated the expression of Reln, Bcl11b and BrdU at both E12.5 and E15. We identified, most Reln+/Bcl11b+ neurons in LII were born at E12.5, however a few were also born later at E15 (Figure 7E). A small number of Reln+/Bcl11b- neurons were born both at E12.5 and E15 (Figure 7E). A small subset of Reln-/Bcl11b+ neurons were born both at E12.5 and at E15 (Figure 7E). This indicates that the most prominent population of neurons born at E12.5 in LII is the Reln+/Bcl11b+ neurons. Together this identifies two Reln+ populations emerging at LII.

## Discussion

We present the boundaries of the MEC and LEC in the developing pig brain and use the pig as a model to investigate neurogenesis and lamination in the MEC. We have found it to be an excellent choice for studying brain development since it closely represents human neurogenesis. We identified that neurogenesis in the pig EC spans from E39 to E60 (during the 1st and early 2nd trimester) which corresponds closely to the timing of neurogenesis in humans (gestational week 7-17) (Kostovic, Sedmak et al. 2019) but is dissimilar to that in mice, in which it extends almost until birth (E11-E17) (Casarosa, Fode et al. 1999, Roy, Kuznicki et al. 2004, Stagni, Giacomini et al. 2015). The anatomical orientation of the pig EC was also similar to that in the developing human brain as we could better view the EC in the coronal plane, similar to the human. In contrast, the EC in rodents is best viewed in the sagittal and horizontal plane. Immunohistochemical analyses revealed that the pig has a moderate OSVZ present from E33-E60 in the EC, which changed in its distribution of intermediate progenitor populations, from a mixed identity of EOMES+/PAX6- and EOMES+/PAX6+ to an enriched population of EOMES+/PAX6+ cells. The entorhinal cells in the pig emerge during the early second trimester and by E60, key cytoarchitectonic features delineate the MEC from the LEC. In humans, the emergence of the EC occurs towards the end of the first trimester with prominent entorhinal cells visible as early as 10.5 weeks post ovulation (Kostovic, Petanjek et al. 1993). The pig, therefore, has a larger OSVZ than rodents and the timing of neurogenesis concurs more closely to neurogenesis in the human. Given it is extremely difficult to obtain human fetal tissue during the second and third trimester, the large gyrencephalic pig brain should be considered a very good model.

Our study revealed novel features within the developing and postnatal EC. We identified pan-neuronal clustering and abundance of OPCs in the LEC and perirhinal cortex early in development at E60 in the superficial layers. The early emergence of these OPCs during gestation suggests that at least one wave of oligodendrogenesis may overlap with EC neurogenesis. In mice, the first OPCs emerge from the ventral medial ganglionic eminence at E11.5 and migrate throughout the telencephalon in a ventral to dorsal manner midway through gestation (Kessaris, Fogarty et al. 2006). Previous research has shown that in the piriform cortex, OPCs are highly plastic and can give rise to pyramidal neurons in the adult mouse (Guo, Maeda et al. 2010). It is not clear what the fate of these OPCS are and whether they have migrated in from the ganglionic eminence or emerged locally by gliogenesis. It is also not clear what the purpose for these cells is in the superficial layers of the LEC and perirhinal cortex or how plastic they may be. We are also uncertain if this is a unique feature in the pig, or if they simply have not received much attention in other species. Further investigation into the role of these clustered OPCs will help to reveal their role and function.

The EC forms part of the periarchicortex and acts as a transitional zone between the allocortex and neocortex. Understanding how it forms may provide insight into the divergence of brain region formation. We unequivocally show that the deep layer neurons are the first neurons to be born in the EC at E10.5 in the mouse. We identified that the MEC forms by parallel lamination events, that differs from classical inside-out lamination events in the neocortex. Similar to the neocortex, the deeper layers precede the birth of the superficial layers, however, both the deeper layers arise in parallel and the superficial layers also arise in parallel. The neurons are scattered evenly across LV/LVI, as well as LII/III. It is not clear what signaling mechanisms may drive this unique parallel lamination patterning in the MEC and whether this is also the case for the LEC and the other periarchicortex regions, such as the presubiculum and parasubiculum. Reln-signaling from the Cajul Retzius cells plays an important role in lamination of the neocortex (D’Arcangelo, Miao et al. 1995, Goldowitz, Cushing et al. 1997) and investigation of how Cajul Retzius cells differ in their signaling in the EC or whether early-born Reln+ neurons contribute to the lamination patterning of the EC would be interesting to follow up in future studies. Ikaros-signaling may also be implicated since it regulates the birth of deep and upper layer neurons in the cerebral cortex (Alsio, Tarchini et al. 2013). Single cell RNA sequencing and analysis studies of excitatory neuron cell fates in the neocortex confirm that deep layer neurons emerge first, prior to upper layer neurons (Loo, Simon et al. 2019, Fan, Fu et al. 2020). Even the recent emergence of flashtag labeling of embryonic progenitors that allows for greater precision in birthdate labeling of neurons (Govindan, Oberst et al. 2018) confirms the classic inside-out lamination pattern of the neocortex (Telley, Agirman et al. 2019). It is plausible to consider that altered lamination patterns in the MEC might relate to altered connectivity and a more thorough investigation of lamination events in the LEC and other periallocortex regions may help to elucidate whether this is a unique feature of the periallocortex.

Studying the emergence of entorhinal cells is interesting since temporal divergences exist in both the emergence and maturation of the cell types in at least the superficial layers. In the mouse, the stellate cells have been shown to emerge in the MEC at approximately E12, with pyramidal neurons emerging approximately 2 days later (Donato, Jacobsen et al. 2017). Our study demonstrates that the early-born entorhinal cells in the superficial layers express Bcl11b+ which is an oddity since Bcl11b is a well-known LV neuron marker in the neocortex. Co-expression of Bcl11b with Reln in the postnatal mouse brain confirms these neurons to be stellate cells. Bcl11b has been shown to be transiently expressed in all deep layer early-born neurons, which is then downregulated in LVI neurons and remains expressed in LV neurons (Kwan, Lam et al. 2008). Furthermore, Bcl11b is a downstream effector of Fezf2 and its expression in neurons in the cerebral cortex mediates subcortical projection of axons (Chen, Wang et al. 2008). However, Reln+ neurons in LII of the MEC project intracortically to the dentate gyrus and CA3 (Kitamura, Pignatelli et al. 2014, Fuchs, Neitz et al. 2016, Leitner, Melzer et al. 2016). We presume therefore that their axonal projections are steered by other signaling mechanisms which may or may not relate to Fezf2 signaling. We also demonstrate in the pig model that the early-born stellate cells are RELN- when they are first deposited in LII and LIII and switch on RELN later. This may have implications for the parallel lamination events, as later-born LII neurons at E14 are able to bypass the LIII neurons and early expression of Reln in the LIII neurons might prevent such migration. It is not clear from our studies whether the early-born stellate cells in mice express Reln at E14, but it would be interesting to investigate this. We demonstrate that the deep layer neurons and the early-born superficial layer neurons arising at E12.5 both express Bcl11b which indicates they arise from a common progenitor. Interestingly, the BrdU labelling in the superficial neurons is weaker than the deep layer neurons and suggests these cells are born one or more cell divisions later. This indicates an important probabilistic decision event occurs in the common progenitor at E12.5. This observation supports the theory of stochastic cortical neurogenesis arising from radial glia progenitors that over time alters the molecular internal clocks within the cells (Llorca, Ciceri et al. 2019). It cannot explain, however, how neuron migration is directed, as the early newborn Reln+ neurons are scattered across both LII and LIII at this time point. Our study clearly shows that in the mouse, LII is the last layer deposited during neurogenesis, which contrasts an earlier study which identified that LIII formed last in the rat (Bayer, Altman et al. 1993). This contrasting finding might relate to species-specific differences and further investigation in a wider range of species is required to conclude whether this is the case. It is important to also investigate lamination patterning in the LEC to form a more meaningful conclusion of general lamination patterns across the EC. We clearly identified subsets of Reln+ cells that differed in their timing of emergence in LII. It is highly likely that the large population of Reln+/Bcl11b+ neurons that were born early on at E12.5 are stellate cells, but the identity of the Reln+/Bcl11b- neurons is not clear. We speculate that these cells correspond to the intermediate stellate/pyramdidal cell described by Fuchs et al., (Fuchs, Neitz et al. 2016) or a second excitatory Reln+ population expressing a unique enhancer from the stellate cells (Blankvoort, Witter et al. 2018). Further investigation is needed to elucidate the identity of the Reln+ cell subtypes we identified in the MEC LII.

To conclude, we characterize neurogenesis events in the developing MEC, in a novel large mammalian model that closely resembles human neurogenesis. We describe for the first time, parallel lamination of the deeper layers followed by parallel lamination of the superficial layers in the MEC which differs from the classic inside-outside lamination of the neocortex. We also identify unique anatomical features in the porcine EC, including prominent clusters of OPCs in the LEC and white matter projections that extend from the EC bilaterally across the brain hemispheres. The EC accelerates markedly in growth at the end of neurogenesis at the end of the second trimester. We highlight particular dynamics in the emergence of Reln+ populations in the LII of the MEC and identify that Reln switches on later in these cells, days after the cells are born. We therefore provide detailed insight into the dynamics of the emerging EC which provides novel insight into altered cortical lamination events compared to the rest of the neocortex. Further studies investigating the mechanisms of parallel lamination may help to determine how this process occurs and whether this is a conserved feature across the periallocortex regions of the brain.

## Author Contributions

YL: Performed immunohistochemistry on the pig brain, DTI/MRI, analyzed data and contributed to the writing of the manuscript

TB: Performed histology of the pig brain, BrdU labeling, analyzed data and contributed to the writing of the manuscript

YM: Performed the DTI/MRI and contributed to the analysis of the DTI data

JMPV: Contributed to immunohistochemistry of pig brain

MP: Contributed to the sectioning and Cresyl violet staining of the pig brain

NAV: Performed BrdU labeling in mice

PDT: Provided porcine fetal specimens

SES: Supervised the project and reviewed the manuscript.

JG: Supervised the project and reviewed the manuscript.

PH: Supervised project and contributed to acquisition of funding and reviewed the manuscript.

KK: Provided materials, personnel and resources for the BrDU labeling and contributed to interpretation of the data

MPW: Contributed to anatomical analysis of the EC in the pig and in acquisition of funding

VH: Contributed to the projects concept, methodology, formal analyses, project’s resources, writing, reviewing & editing the manuscript. Administered and managed the project and acquired the project’s funding.

All authors read and approved the final manuscript

## Acknowledgments

We thank Per Torp Sangild for his donation of healthy adult sow brains for this study. We acknowledge the Core Facility for Integrated Microscopy, Faculty of Health and Medical Sciences, University of Copenhagen for access use of the Axio Scan Z1.

## Data Availability Statement

No datasets available in this study

---

**Figure S1. Borders of the developing porcine entorhinal cortex (EC).** Cresyl violet stained 4 or 5 coronal sections of the piriform lobe from Embryonic day (E)60 to postnatal day (P)75 depicted in a rostral to caudal series. The first section is rostral to the EC, the second section depicts the LEC occupying the EC entity, the third section includes both MEC and LEC present, the fourth section depicts the MEC occupying the entire mediolateral entity and the fifth section is the most caudal part of the piriform lobe. Dentate gyrus (DG); hippocampal area (HA); amygdala (Amyg); posterior rhinal sulcus (RHP); medial entorhinal cortex (MEC); lateral entorhinal cortex (LEC); pre-subiculum (PreS); para-subiculum (PaS), subiculum (Sub); perirhinal cortex (PER). Scale bar 1 cm.

**Figure S2. The morphology of the entorhinal cortex at Embryonic day (E)50 to E70.** Cresyl violet staining of the developing cortex show a prominent layer or entorhinal neurons with large nuclei in the superficial layer from E50 onwards whereas, the glia cells are difficult to identify at E50 from the nissl staining alone. Scale bar 200 μm.

**Figure S3 Characterization of the germinal layers in the developing porcine entorhinal cortex (EC). (A)** A schematic overview of the location of the characterized medial EC (MEC). Axes: D, dorsal; V, Ventral; A, Anterior; P, posterior **(B)** Expression of GFAP and FABP7 in the MEC ventricular zone (VZ). Scale bar 25 μm. **(C)** Temporal expression of radial glia (GFAP, FABP7, PAX6, SOX2) during MEC development. Scale bar 50 μm **(D)**. Quantification of the thickness in μm of the VZ during development. **(E)** Expression of EOMES and PAX6 in the EC. Scale bar 25 μm (up) and 50μm (bottom). **(F)** Quantification of the EOMES+/PAX6+ and EOMES+/PAX6-cell populations in the germinal zone. **(G)** TBR1/EOMES/RELN expression during development. Scale bar 50 μm. **(H)** Expression of TBR1/EOMES/RELN in the marginal zone and the cortical plate, enlarged from red boxes in G **(F)**. Scale bar 25 μm. (HO = Hoeschst, V = ventral telencephalon). Error bars represent SD.

**Figure S4. Comparative expression of superficial and deep layer markers in the early developing entorhinal cortex (EC) versus the dorsal, cingulate gyrus.** Expression of the canonical deep layer marker (BCL11B), superficial layer marker (SATB2) and stellate cell / Cajal–Retzius cells (CR cells) marker RELN from Embryonic day (E)26 to E39 shows the prevalence of BCL11B and SATB2 from E33 onwards in the superficial marginalzone/cortical plate and the absence of RELN at these time points within the developing cortical plate. Scale bar 50 μm.

**Figure S5. Laminar birth dating of the medial entorhinal cortex (MEC) in the mouse.** Bromodeoxyuri-dine (BrdU) labelling in combination with analysis of NeuN expression of postnatal day (P)7 MEC shows the emergence of newborn neurons from embryonic day (E)10 to E16. **(A)** Representative z-stack images from BrdU injections from E9-E16. Dotted lines represent the LII/LIII boundary and the LIV *lamina dissecans*. Insert boxes are shown at higher magnification in B. White arrowheads depict cells highlighted in B. Scale bar 25 μm. **(B)** High magnification of representative BrdU labelled cells with overlapping expression of NeuN in a single z-plane from LII and LIII areas in A. White arrows highlight BrdU labeled cells. Yellow asterix (*) denotes cells with strong BrdU labelling which were considered to be born at the time of BrdU injection. Scale bar 10 μm.

**Supplementary Table 1.** Connectivity of neuronal tracts to/from the lateral entorhinal cortex (LEC) and the medial entorhinal cortex (MEC) in the postnatal pig (P75). The average connectivity from three analyzed brains is shown with the standard deviation in brackets. Abbreviations: AM, amygdala; CN, caudate nucleus, DG, dentate gyrus; HC, dorsal hippocampal commissure; LEC, lateral entorhinal cortex; LOT, lateral olfactory tract; MEC, medial entorhinal cortex; Nucleus accumbens, NAc; PUT, putamen; SUB, subiculum.

|  | Total Tracts | MEC | LEC | AM | CA1 | CA3 | DG | PUT | SUB | LOT | CN | NAc | HC | Other |
| --- | --- | --- | --- | --- | --- | --- | --- | --- | --- | --- | --- | --- | --- | --- |
| **MEC** | 6205 |  | 106<br>(1.71) | 730.3<br>(11.77) | 91.3<br>(1.47) | 49.3<br>(0.79) | 47<br>(0.76) | 4.7<br>(0.08) | 534<br>(8.61) | 3.3<br>(0.05) | 9.0<br>(0.15) | 1.0<br>(0.02) | 11.7<br>(0.19) | 4617.3<br>(74.41) |
| **LEC** | 4516.3 | 106<br>(2.35) |  | 468.7<br>(10.38) | 5.3<br>(0.12) | 10<br>(0.22) | 2.7<br>(0.06) | 59<br>(1.31) | 1<br>(0.02) | 14.3<br>(0.32) | 75.3<br>(1.67) | 0<br>(0) | 0<br>(0) | 3774<br>(83.56) |

## References

Alsio, J. M., B. Tarchini, M. Cayouette and F. J. Livesey (2013). “Ikaros promotes early-born neuronal fates in the cerebral cortex”. Proc Natl Acad Sci U S A 110(8): E716–725.

Bayer, S. A. (1980). “Development of the hippocampal region in the rat. I. Neurogenesis examined with 3H-thymidine autoradiography”. J Comp Neurol 190(1): 87–114.

Bayer, S. A. (1980). “Development of the hippocampal region in the rat. II. Morphogenesis during embryonic and early postnatal life”. J Comp Neurol 190(1): 115–134.

Bayer, S. A., J. Altman, R. J. Russo and X. Zhang (1993). “Timetables of neurogenesis in the human brain based on experimentally determined patterns in the rat”. Neurotoxicology 14(1): 83–144.

Bergmann, E., G. Zur, G. Bershadsky and I. Kahn (2016). “The Organization of Mouse and Human Cortico-Hippocampal Networks Estimated by Intrinsic Functional Connectivity”. Cereb Cortex 26(12): 4497–4512.

Bjarkam, C. R., A. N. Glud, D. Orlowski, J. C. H. Sorensen and N. Palomero-Gallagher (2017). “The telencephalon of the Gottingen minipig, cytoarchitecture and cortical surface anatomy”. Brain Struct Funct 222(5): 2093–2114.

Blankvoort, S., M. P. Witter, J. Noonan, J. Cotney and C. Kentros (2018). “Marked Diversity of Unique Cortical Enhancers Enables Neuron-Specific Tools by Enhancer-Driven Gene Expression”. Curr Biol 28(13): 2103–2114 e2105.

Burwell, R. D. and D. G. Amaral (1998). “Perirhinal and postrhinal cortices of the rat: interconnectivity and connections with the entorhinal cortex”. J Comp Neurol 391(3): 293–321.

Canto, C. B. and M. P. Witter (2012). “Cellular properties of principal neurons in the rat entorhinal cortex. I. The lateral entorhinal cortex”. Hippocampus 22(6): 1256–1276.

Canto, C. B. and M. P. Witter (2012). “Cellular properties of principal neurons in the rat entorhinal cortex. II. The medial entorhinal cortex”. Hippocampus 22(6): 1277–1299.

Casarosa, S., C. Fode and F. Guillemot (1999). “Mash1 regulates neurogenesis in the ventral telencephalon”. Development 126(3): 525–534.

Chen, B., S. S. Wang, A. M. Hattox, H. Rayburn, S. B. Nelson and S. K. McConnell (2008). “The Fezf2-Ctip2 genetic pathway regulates the fate choice of subcortical projection neurons in the developing cerebral cortex”. Proc Natl Acad Sci U S A 105(32): 11382–11387.

Conrad, M. S., R. N. Dilger and R. W. Johnson (2012). “Brain growth of the domestic pig (Sus scrofa) from 2 to 24 weeks of age: a longitudinal MRI study”. Dev Neurosci 34(4): 291–298.

Conrad, M. S., B. P. Sutton, R. N. Dilger and R. W. Johnson (2014). “An in vivo three-dimensional magnetic resonance imaging-based averaged brain collection of the neonatal piglet (Sus scrofa)”. PLoS One 9(9): e107650.

Cooper, J. A. (2008). “A mechanism for inside-out lamination in the neocortex”. Trends Neurosci 31(3): 113–119.

D’Arcangelo, G., G. G. Miao, S. C. Chen, H. D. Soares, J. I. Morgan and T. Curran (1995). “A protein related to extracellular matrix proteins deleted in the mouse mutant reeler”. Nature 374(6524): 719–723.

Domnisoru, C., A. A. Kinkhabwala and D. W. Tank (2013). “Membrane potential dynamics of grid cells”. Nature 495(7440): 199–204.

Donato, F., R. I. Jacobsen, M. B. Moser and E. I. Moser (2017). “Stellate cells drive maturation of the entorhinal-hippocampal circuit”. Science 355(6330).

Fan, X., Y. Fu, X. Zhou, L. Sun, M. Yang, M. Wang, R. Chen, Q. Wu, J. Yong, J. Dong, L. Wen, J. Qiao, X. Wang and F. Tang (2020). “Single-cell transcriptome analysis reveals cell lineage specification in temporal-spatial patterns in human cortical development”. Sci Adv 6(34): eaaz2978.

Farr, M., G. D. Kitas, L. Waterhouse, R. Jubb, D. Felix-Davies and P. A. Bacon (1988). “Treatment of psoriatic arthritis with sulphasalazine: a one year open study”. Clin Rheumatol 7(3): 372–377.

Fietz, S. A., I. Kelava, J. Vogt, M. Wilsch-Brauninger, D. Stenzel, J. L. Fish, D. Corbeil, A. Riehn, W. Distler, R. Nitsch and W. B. Huttner (2010). “OSVZ progenitors of human and ferret neocortex are epithelial-like and expand by integrin signaling”. Nat Neurosci 13(6): 690–699.

Fuchs, E. C., A. Neitz, R. Pinna, S. Melzer, A. Caputi and H. Monyer (2016). “Local and Distant Input Controlling Excitation in Layer II of the Medial Entorhinal Cortex”. Neuron 89(1): 194–208.

Garcia-Cabezas, M. A., Y. J. John, H. Barbas and B. Zikopoulos (2016). “Distinction of Neurons, Glia and Endothelial Cells in the Cerebral Cortex: An Algorithm Based on Cytological Features”. Front Neuroanat 10: 107.

Gatome, C. W., L. Slomianka, H. P. Lipp and I. Amrein (2010). “Number estimates of neuronal phenotypes in layer II of the medial entorhinal cortex of rat and mouse”. Neuroscience 170(1): 156–165.

Goldowitz, D., R. C. Cushing, E. Laywell, G. D’Arcangelo, M. Sheldon, H. O. Sweet, M. Davisson, D. Steindler and T. Curran (1997). “Cerebellar disorganization characteristic of reeler in scrambler mutant mice despite presence of reelin”. J Neurosci 17(22): 8767–8777.

Govindan, S., P. Oberst and D. Jabaudon (2018). “In vivo pulse labeling of isochronic cohorts of cells in the central nervous system using FlashTag”. Nat Protoc 13(10): 2297–2311.

Guo, F., Y. Maeda, J. Ma, J. Xu, M. Horiuchi, L. Miers, F. Vaccarino and D. Pleasure (2010). “Pyramidal neurons are generated from oligodendroglial progenitor cells in adult piriform cortex”. J Neurosci 30(36): 12036–12049.

Hevner, R. F., T. Neogi, C. Englund, R. A. Daza and A. Fink (2003). “Cajal-Retzius cells in the mouse: transcription factors, neurotransmitters, and birthdays suggest a pallial origin”. Brain Res Dev Brain Res 141(1-2): 39–53.

Holm, I. E. and F. A. Geneser (1989). “Histochemical demonstration of zinc in the hippocampal region of the domestic pig: I. Entorhinal area, parasubiculum, and presubiculum”. J Comp Neurol 287(2): 145–163.

Insausti, R., M. T. Herrero and M. P. Witter (1997). “Entorhinal cortex of the rat: cytoarchitectonic subdivisions and the origin and distribution of cortical efferents”. Hippocampus 7(2): 146–183.

Insausti, R., M. Munoz-Lopez, A. M. Insausti and E. Artacho-Perula (2017). “The Human Periallocortex: Layer Pattern in Presubiculum, Parasubiculum and Entorhinal Cortex. A Review”. Front Neuroanat 11: 84.

Insausti, R., T. Tunon, T. Sobreviela, A. M. Insausti and L. M. Gonzalo (1995). “The human entorhinal cortex: a cytoarchitectonic analysis”. J Comp Neurol 355(2): 171–198.

Jelsing, J., R. Nielsen, A. K. Olsen, N. Grand, R. Hemmingsen and B. Pakkenberg (2006). “The postnatal development of neocortical neurons and glial cells in the Gottingen minipig and the domestic pig brain”. J Exp Biol 209(Pt 8): 1454–1462.

Kessaris, N., M. Fogarty, P. Iannarelli, M. Grist, M. Wegner and W. D. Richardson (2006). “Competing waves of oligodendrocytes in the forebrain and postnatal elimination of an embryonic lineage”. Nat Neurosci 9(2): 173–179.

Kim, S. W. (2010). “Recent advances in sow nutrition”. Revista Brasileira de Zootecnia 39: 303–310.

Kitamura, T., M. Pignatelli, J. Suh, K. Kohara, A. Yoshiki, K. Abe and S. Tonegawa (2014). “Island cells control temporal association memory”. Science 343(6173): 896–901.

Kolenkiewicz, M., A. Robak, M. Rowniak, K. Bogus-Nowakowska, J. Calka and M. Majewski (2009). “Distribution of cocaine- and amphetamine-regulated transcript in the hippocampal formation of the guinea pig and domestic pig”. Folia Morphol (Warsz) 68(1): 23–31.

Kostovic, I., Z. Petanjek and M. Judas (1993). “Early areal differentiation of the human cerebral cortex: entorhinal area”. Hippocampus 3(4): 447–458.

Kostovic, I., G. Sedmak and M. Judas (2019). “Neural histology and neurogenesis of the human fetal and infant brain”. Neuroimage 188: 743–773.

Kwan, K. Y., M. M. Lam, Z. Krsnik, Y. I. Kawasawa, V. Lefebvre and N. Sestan (2008). “SOX5 postmitotically regulates migration, postmigratory differentiation, and projections of subplate and deep-layer neocortical neurons”. Proc Natl Acad Sci U S A 105(41): 16021–16026.

Leitner, F. C., S. Melzer, H. Lutcke, R. Pinna, P. H. Seeburg, F. Helmchen and H. Monyer (2016). “Spatially segregated feedforward and feedback neurons support differential odor processing in the lateral entorhinal cortex”. Nat Neurosci 19(7): 935–944.

Llorca, A., G. Ciceri, R. Beattie, F. K. Wong, G. Diana, E. Serafeimidou-Pouliou, M. Fernandez-Otero, C. Streicher, S. J. Arnold, M. Meyer S. Hippenmeyer, M. Maravall and O. Marin (2019). “A stochastic framework of neurogenesis underlies the assembly of neocortical cytoarchitecture”. Elife 8.

Loo, L., J. M. Simon, L. Xing, E. S. McCoy, J. K. Niehaus, J. Guo, E. S. Anton and M. J. Zylka (2019). “Single-cell transcriptomic analysis of mouse neocortical development”. Nat Commun 10(1): 134.

Martinez-Cerdeno, V., C. L. Cunningham, J. Camacho, J. L. Antczak, A. N. Prakash, M. E. Cziep, A. I. Walker and S. C. Noctor (2012). “Comparative analysis of the subventricular zone in rat, ferret and macaque: evidence for an outer subventricular zone in rodents”. PLoS One 7(1): e30178.

Mufson, E. J., D. R. Brady and J. H. Kordower (1990). “Tracing neuronal connections in postmortem human hippocampal complex with the carbocyanine dye DiI”. Neurobiol Aging 11(6): 649–653.

Patten, A. R., C. J. Fontaine and B. R. Christie (2014). “A comparison of the different animal models of fetal alcohol spectrum disorders and their use in studying complex behaviors”. Front Pediatr 2: 93.

Pontelo, T. P., J. R. Miranda, M. A. R. Felix, B. A. Pereira, W. E. da Silva, G. F. Avelar, F. Mariano, G. C. Guimaraes and M. G. Zangeronimo (2018). “Histological characteristics of the gonads of pig fetuses and their relationship with fetal anatomical measurements”. Res Vet Sci 117: 28–36.

Ray, S. and M. Brecht (2016). “Structural development and dorsoventral maturation of the medial entorhinal cortex”. Elife 5: e13343.

Rowland, D. C., H. A. Obenhaus, E. R. Skytoen, Q. Zhang, C. G. Kentros, E. I. Moser and M. B. Moser (2018). “Functional properties of stellate cells in medial entorhinal cortex layer II”. Elife 7: e36664.

Rowland, D. C., A. P. Weible, I. R. Wickersham, H. Wu M. Mayford, M. P. Witter and C. G. Kentros (2013). “Transgenically targeted rabies virus demonstrates a major monosynaptic projection from hippocampal area CA2 to medial entorhinal layer II neurons”. J Neurosci 33(37): 14889–14898.

Roy, K., K. Kuznicki, Q. Wu, Z. Sun, D. Bock, G. Schutz, N. Vranich and A. P. Monaghan (2004). “The Tlx gene regulates the timing of neurogenesis in the cortex”. J Neurosci 24(38): 8333–8345.

Saikali, S., P. Meurice, P. Sauleau, P. A. Eliat, P. Bellaud, G. Randuineau, M. Verin and C. H. Malbert (2010). “A three-dimensional digital segmented and deformable brain atlas of the domestic pig”. J Neurosci Methods 192(1): 102–109.

Sansom, S. N., D. S. Griffiths, A. Faedo, D. J. Kleinjan, Y. Ruan, J. Smith, V. van Heyningen, J. L. Rubenstein and F. J. Livesey (2009). “The level of the transcription factor Pax6 is essential for controlling the balance between neural stem cell self-renewal and neurogenesis”. PLoS Genet 5(6): e1000511.

Schmidt-Hieber, C. and M. Hausser (2013). “Cellular mechanisms of spatial navigation in the medial entorhinal cortex”. Nat Neurosci 16(3): 325–331.

Schultz, H., T. Sommer and J. Peters (2015). “The Role of the Human Entorhinal Cortex in a Representational Account of Memory”. Front Hum Neurosci 9: 628.

Smart, I. H., C. Dehay, P. Giroud, M. Berland and H. Kennedy (2002). “Unique morphological features of the proliferative zones and postmitotic compartments of the neural epithelium giving rise to striate and extrastriate cortex in the monkey”. Cereb Cortex 12(1): 37–53.

Stagni, F., A. Giacomini, S. Guidi, E. Ciani and R. Bartesaghi (2015). “Timing of therapies for Down syndrome: the sooner, the better”. Front Behav Neurosci 9: 265.

Staubli, U., G. Ivy and G. Lynch (1984). “Hippocampal denervation causes rapid forgetting of olfactory information in rats”. Proc Natl Acad Sci U S A 81(18): 5885–5887.

Stephan, H. (1975). Allocortex. Handbuch der mikroskopischen Anatomie des Menschen. W. H. Bargmann. Bd 4/9, Berlin Heidelberg, New York, Springer: S1–998.

Stephan, H. (1983). “Evolutionary trend in limbic structures”. Neurosci Behav Rev 7: 367–374.

Stephan, H. and O. J. Andy (1970). The allocortex in primates. Advances in Primatology. C. R. Noback and W. Montagna. New York, Appleton Century Croft: 289–297.

Sun, Y., A. Q. Nguyen, J. P. Nguyen, L. Le, D. Saur, J. Choi, E. M. Callaway and X. Xu (2014). “Cell-type-specific circuit connectivity of hippocampal CA1 revealed through Cre-dependent rabies tracing”. Cell Rep 7(1): 269–280.

Telley, L., G. Agirman, J. Prados, N. Amberg, S. Fievre, P. Oberst, G. Bartolini, I. Vitali, C. Cadilhac, S. Hippenmeyer, L. Nguyen, A. Dayer and D. Jabaudon (2019). “Temporal patterning of apical progenitors and their daughter neurons in the developing neocortex”. Science 364(6440).

Totterdell, S. and G. E. Meredith (1997). “Topographical organization of projections from the entorhinal cortex to the striatum of the rat”. Neuroscience 78(3): 715–729.

Vandrey, B., D. L. F. Garden, V. Ambrozova, C. McClure, M. F. Nolan and J. A. Ainge (2020). “Fan Cells in Layer 2 of the Lateral Entorhinal Cortex Are Critical for Episodic-like Memory”. Curr Biol 30(1): 169–175 e165.

Vasistha, N. A., M. Johnstone, S. K. Barton, S. E. Mayerl, B. Thangaraj Selvaraj, P. A. Thomson, O. Dando, E. Grunewald, C. Alloza, M. E. Bastin, M. R. Livesey, K. Economides, D. Magnani, P. Makedonopolou, K. Burr, D. J. Story, D. H. R. Blackwood, D. J. A. Wyllie, A. M. McIntosh, J. K. Millar, C. Ffrench-Constant, G. E. Hardingham, S. M. Lawrie and S. Chandran (2019). “Familial t(1;11) translocation is associated with disruption of white matter structural integrity and oligodendrocyte-myelin dysfunction”. Mol Psychiatry 24(11): 1641–1654.

Wilhite, B. L., T. J. Teyler and C. Hendricks (1986). “Functional relations of the rodent claustral-entorhinal-hippocampal system”. Brain Res 365(1): 54–60.

Winter, J. D., S. Dorner, J. Lukovic, J. A. Fisher, K. S. St Lawrence and A. Kassner (2011). “Noninvasive MRI measures of microstructural and cerebrovascular changes during normal swine brain development”. Pediatr Res 69(5 Pt 1): 418–424.

Witter, M. P., T. P. Doan, B. Jacobsen, E. S. Nilssen and S. Ohara (2017). “Architecture of the Entorhinal Cortex A Review of Entorhinal Anatomy in Rodents with Some Comparative Notes”. Front Syst Neurosci 11: 46.

Witter, M. P., P. Room, H. J. Groenewegen and A. H. Lohman (1986). “Connections of the parahippocampal cortex in the cat. V. Intrinsic connections; comments on input/output connections with the hippocampus”. J Comp Neurol 252(1): 78–94.

Wouterlood, F. G., W. Hartig, G. Bruckner and M. P. Witter (1995). “Parvalbumin-immunoreactive neurons in the entorhinal cortex of the rat: localization, morphology, connectivity and ultrastructure”. J Neurocytol 24(2): 135–153.

Woznicka, A., M. Malinowska and A. Kosmal (2006). “Cytoarchitectonic organization of the entorhinal cortex of the canine brain”. Brain Res Rev 52(2): 346–367.

Wyss, J. M., B. Sripanidkulchai and T. L. Hickey (1983). “An analysis of the time of origin of neurons in the entorhinal and subicular cortices of the cat”. J Comp Neurol 221(3): 341–357.

Yushkevich, P. A., J. Piven, H. C. Hazlett, R. G. Smith, S. Ho, J. C. Gee and G. Gerig (2006). “User-guided 3D active contour segmentation of anatomical structures: significantly improved efficiency and reliability”. Neuroimage 31(3): 1116–1128.

Zykin, P. A., I. A. Moiseenko, L. A. Tkachenko, R. A. Nasyrov, E. A. Tsvetkov and E. I. Krasnoshchekova (2018). “Peculiarities of Cyto- and Chemoarchitectonics of Human Entorhinal Cortex during the Fetal Period”. Bull Exp Biol Med 164(4): 497–501.

