## Supplementary material for "Development of the entorhinal cortex occurs via parallel lamination during neurogenesis": Supp Figure 1 and 2

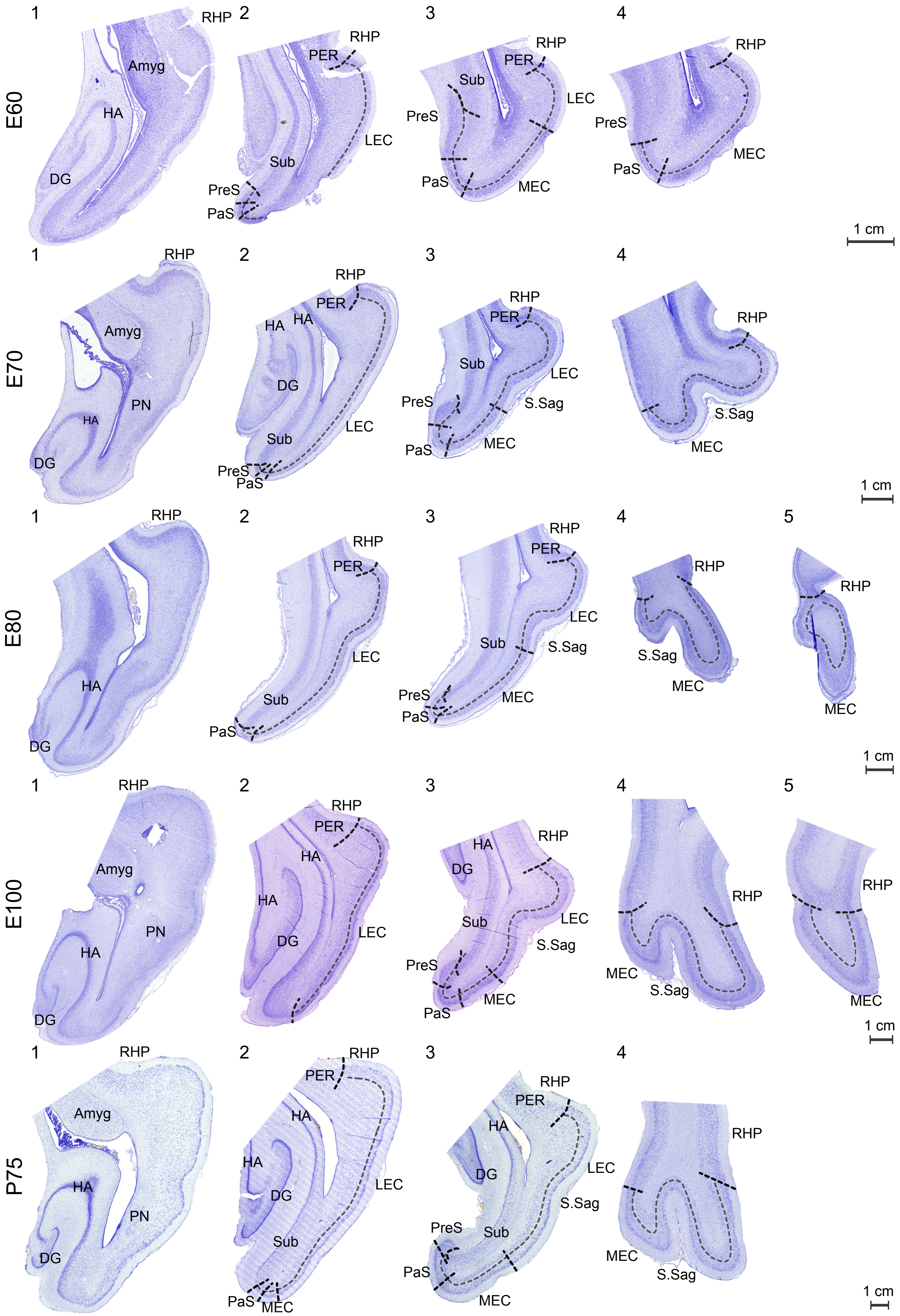
E60
1
RHP
Amyg
HA
DG
2
RHP
PER
LEC
Sub
PreS
PaS
3
RHP
PER
Sub
LEC
PreS
PaS
MEC
4
RHP
PreS
PaS
MEC
1 cm
E70
1
RHP
Amyg
HA
PN
DG
2
RHP
PER
HA
HA
DG
LEC
Sub
PreS
PaS
3
RHP
PER
Sub
LEC
PreS
S.Sag
PaS
MEC
4
RHP
S.Sag
MEC
1 cm
E80
1
RHP
HA
DG
2
RHP
PER
LEC
Sub
PaS
3
RHP
PER
LEC
Sub
S.Sag
PreS
PaS
MEC
4
RHP
S.Sag
MEC
5
RHP
MEC
1 cm
E100
1
RHP
Amyg
PN
HA
DG
2
RHP
PER
HA
HA
LEC
DG
3
HA
RHP
DG
Sub
LEC
S.Sag
PreS
PaS
MEC
4
RHP
MEC
S.Sag
5
RHP
MEC
1 cm
P75
1
RHP
Amyg
HA
PN
DG
2
RHP
PER
HA
HA
DG
LEC
Sub
PaS
MEC
3
RHP
PER
HA
DG
LEC
S.Sag
PreS
Sub
PaS
MEC
4
RHP
MEC
S.Sag
1 cm

**Figure S1. Borders of the developing porcine entorhinal cortex (EC).** Cresyl violet stained 4 or 5 coronal sections of the piriform lobe from Embryonic day (E)60 to postnatal day (P)75 depicted in a rostral to caudal series. The first section is rostral to the EC, the second section depicts the LEC occupying the EC entity, the third section includes both MEC and LEC present, the fourth section depicts the MEC occupying the entire mediolateral entity and the fifth section is the most caudal part of the piriform lobe. Dentate gyrus (DG); hippocampal area (HA); amygdala (Amyg); posterior rhinal sulcus (RHP); medial entorhinal cortex (MEC); lateral entorhinal cortex (LEC); pre-subiculum (PreS); para-subiculum (PAS), subiculum (Sub); perirhinal cortex (PER). Scale bar 1 cm.

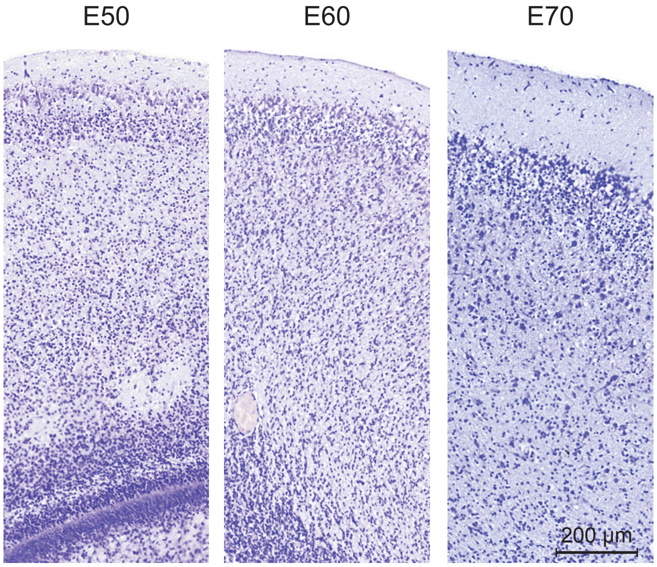


**Figure S2. The morphology of the entorhinal cortex at Embryonic day (E)50 to E70.** Cresyl violet staining of the developing cortex show a prominent layer or entorhinal neurons with large nuclei in the superficial layer from E50 onwards whereas, the glia cells are difficult to identify at E50 from the nissl staining alone. Scale bar 200 µm.
