## Supplementary material for "Development of the entorhinal cortex occurs via parallel lamination during neurogenesis": Supp Figure 3

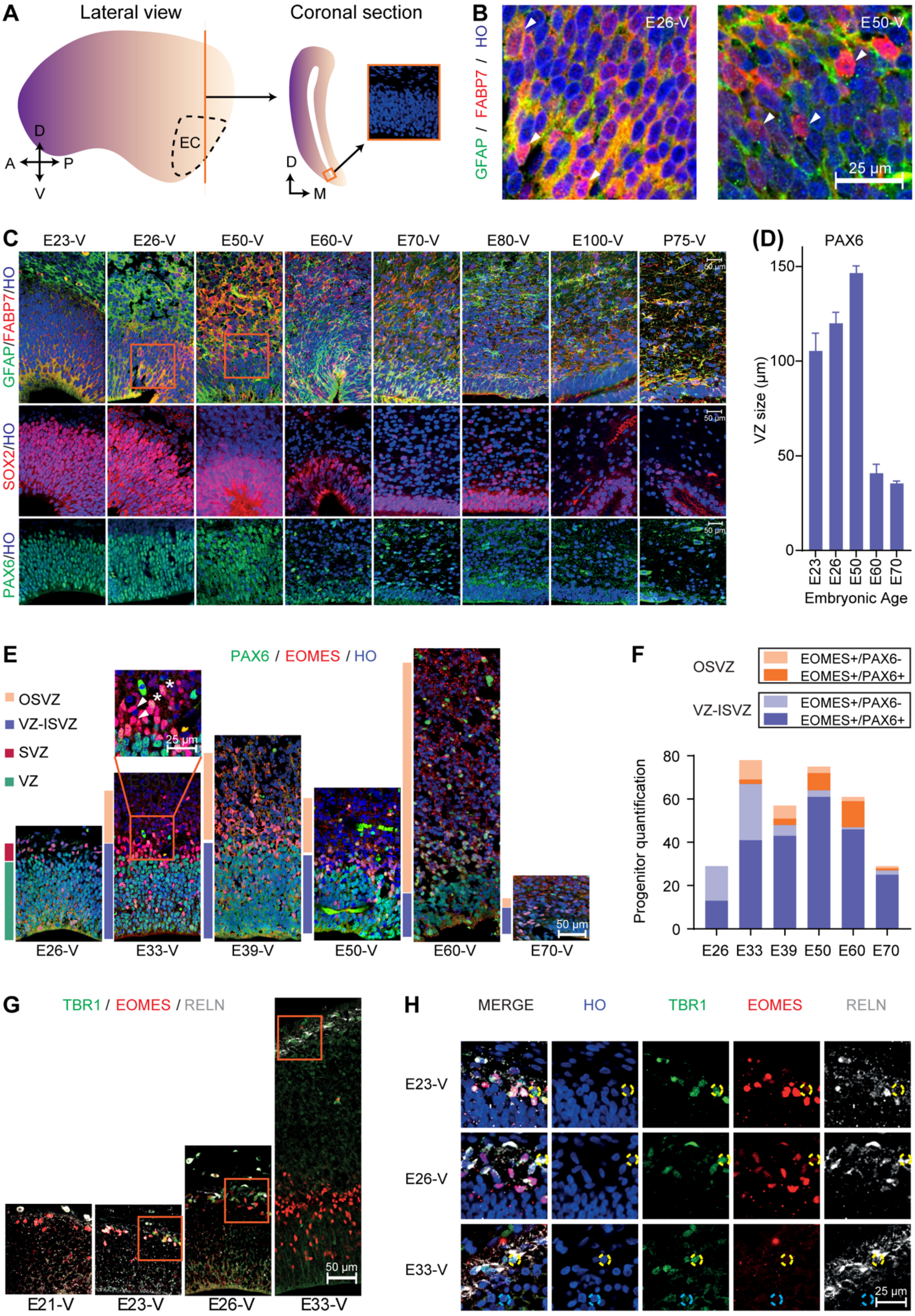
A
Lateral view
Coronal section
EC
D
A
P
V
D
M
B
GFAP / FABP7 / HO
E26-V
E50-V
25 µm
C
E23-V
E26-V
E50-V
E60-V
E70-V
E80-V
E100-V
P75-V
GFAP/FABP7/HO
SOX2/HO
PAX6/HO
50 µm
(D)
PAX6
VZ size (µm)
150
100
50
0
E23
E26
E50
E60
E70
Embryonic Age
E
PAX6 / EOMES / HO
OSVZ
VZ-ISVZ
SVZ
VZ
25 µm
50 µm
E26-V
E33-V
E39-V
E50-V
E60-V
E70-V
F
OSVZ
EOMES+/PAX6-
EOMES+/PAX6+
VZ-ISVZ
EOMES+/PAX6-
EOMES+/PAX6+
Progenitor quantification
80
60
40
20
0
E26
E33
E39
E50
E60
E70
G
TBR1 / EOMES / RELN
50 µm
E21-V
E23-V
E26-V
E33-V
H
MERGE
HO
TBR1
EOMES
RELN
E23-V
E26-V
E33-V
25 µm

**Figure S3 Characterization of the germinal layers in the developing porcine entorhinal cortex (EC). (A)** A schematic overview of the location of the characterized medial EC (MEC). Axes: D, dorsal; V, Ventral; A, Anterior; P, posterior **(B)** Expression of GFAP and FABP7 in the MEC ventricular zone (VZ). Scale bar 25 µm. **(C)** Temporal expression of radial glia (GFAP, FABP7, PAX6, SOX2) during MEC development. Scale bar 50 µm **(D)**. Quantification of the thickness in µm of the VZ during development. **(E)** Expression of EOMES and PAX6 in the EC. Scale bar 25 µm (up) and 50µm (bottom). **(F)** Quantification of the EOMES+/PAX6+ and EOMES+/PAX6- cell populations in the germinal zone. **(G)** TBR1/EOMES/RELN expression during development. Scale bar 50 µm. **(H)** Expression of TBR1/EOMES/RELN in the marginal zone and the cortical plate, enlarged from red boxes in G **(F)**. Scale bar 25 µm. (HO = Hoeschst, V = ventral telencephalon). Error bars represent SD.
