## Supplementary material for "Development of the entorhinal cortex occurs via parallel lamination during neurogenesis": Supp Figure 4

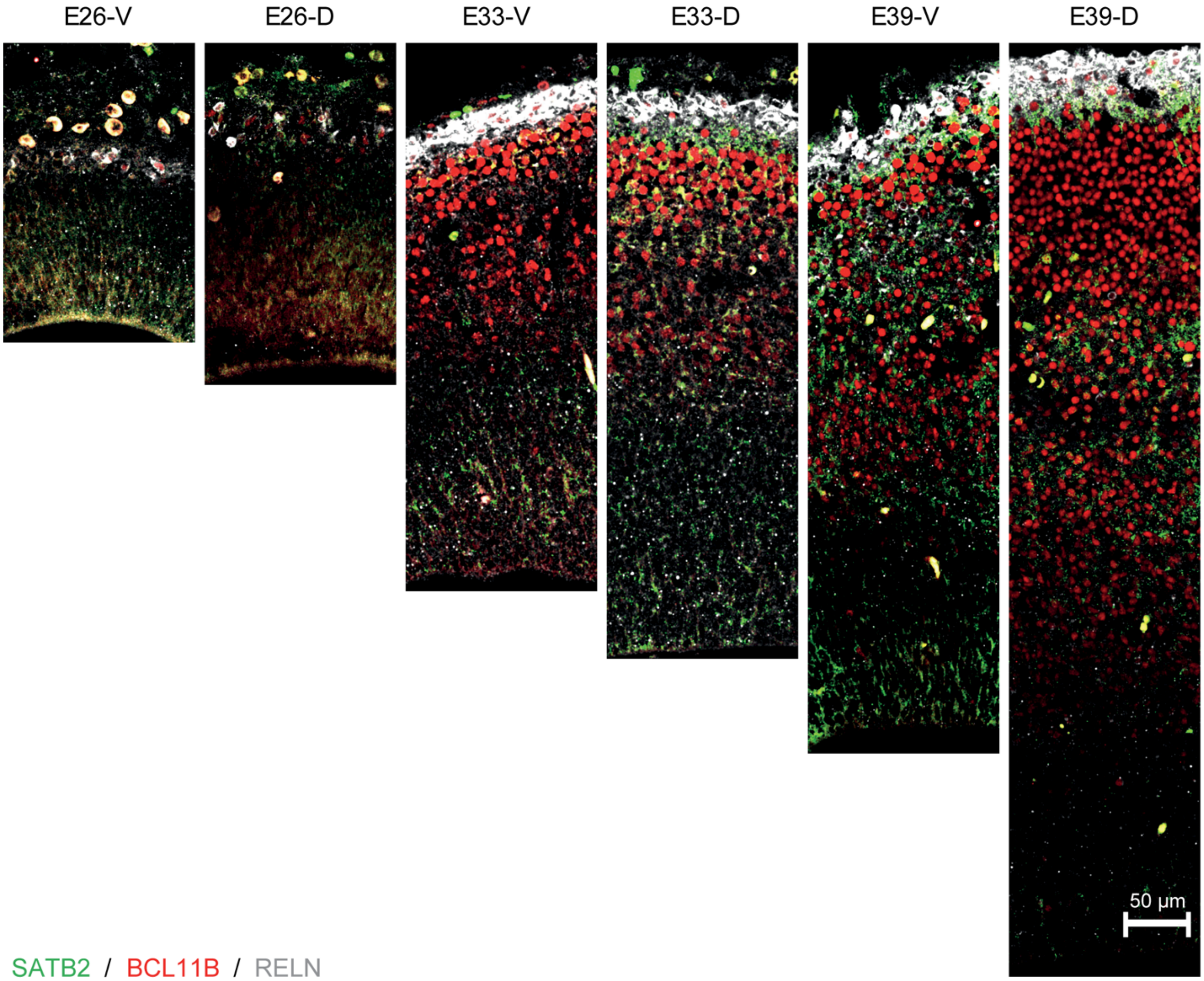


**Figure S4. Comparative expression of superficial and deep layer markers in the early developing entorhinal cortex (EC) versus the dorsal, cingulate gyrus.** Expression of the canonical deep layer marker (BCL11B), superficial layer marker (SATB2) and stellate cell / Cajal–Retzius cells (CR cells) marker RELN from Embryonic day (E)26 to E39 shows the prevalence of BCL11B and SATB2 from E33 onwards in the superficial marginalzone/cortical plate and the absence of RELN at these time points within the developing cortical plate. Scale bar 50 µm.
