## Supplementary material for "Development of the entorhinal cortex occurs via parallel lamination during neurogenesis": Supp Figure 5

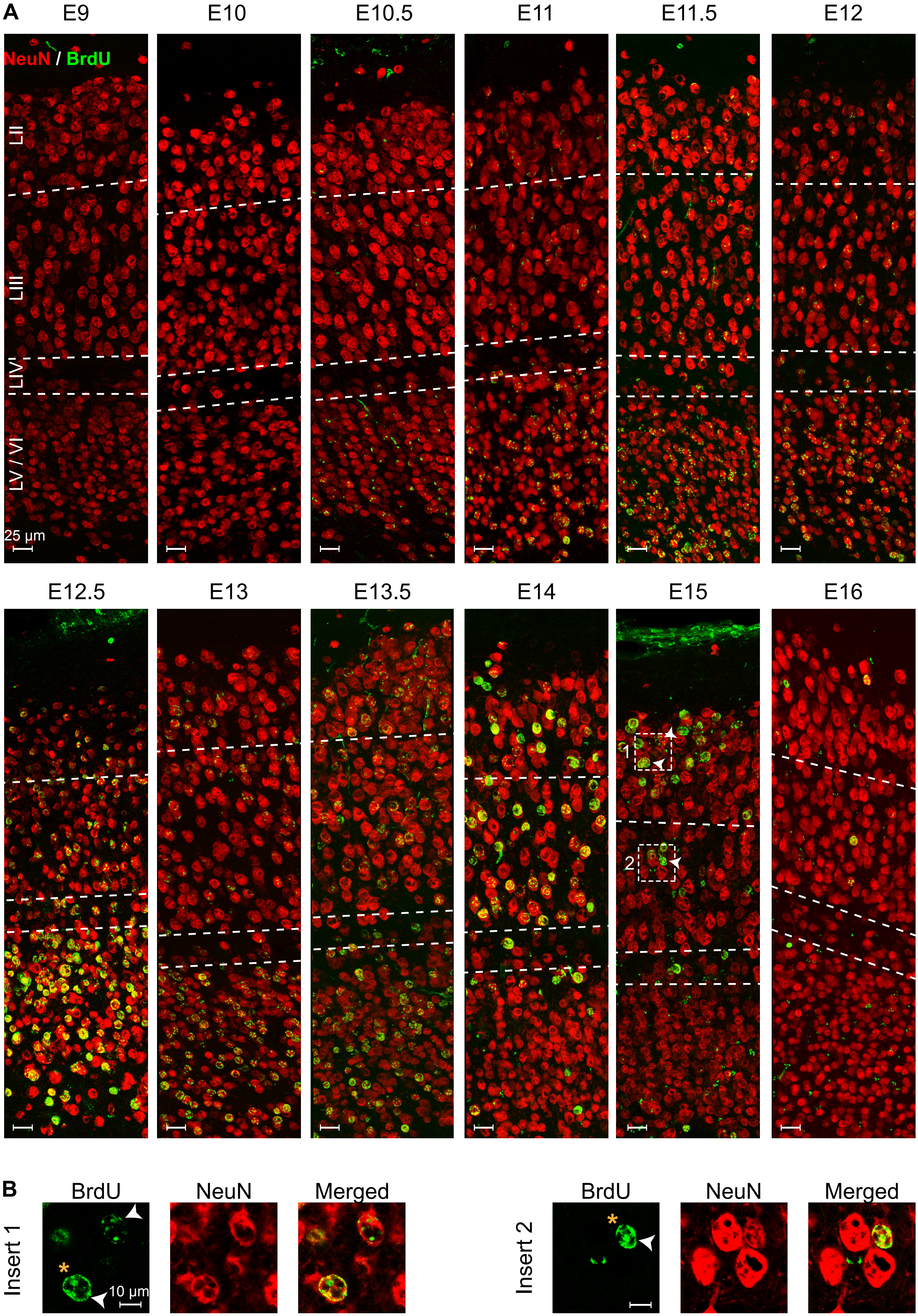
A
E9
E10
E10.5
E11
E11.5
E12
NeuN / BrdU
LII
LIII
LIV
LV / VI
25 µm
E12.5
E13
E13.5
E14
E15
E16
1
2
B
BrdU
NeuN
Merged
Insert 1
10 µm
BrdU
NeuN
Merged
Insert 2

**Figure S5. Laminar birth dating of the medial entorhinal cortex (MEC) in the mouse.** Bromodeoxyuri-dine (BrdU) labelling in combination with analysis of NeuN expression of postnatal day (P)7 MEC shows the emergence of newborn neurons from embryonic day (E)10 to E16. **(A)** Representative z-stack images from BrdU injections from E9-E16. Dotted lines represent the LII/LIII boundary and the LIV *lamina dissecans*. Insert boxes are shown at higher magnification in B. White arrowheads depict cells highlighted in B. Scale bar 25 μm. **(B)** High magnification of representative BrdU labelled cells with overlapping expression of NeuN in a single z-plane from LII and LIII areas in A. White arrows highlight BrdU labeled cells. Yellow asterix (*) denotes cells with strong BrdU labelling which were considered to be born at the time of BrdU injection. Scale bar 10 μm.
